# Abundant Parent-of-origin Effect eQTL: The Framingham Heart Study

**DOI:** 10.1101/2024.06.05.597677

**Authors:** Yongtao Guan, Tianxiao Huan, Daniel Levy

## Abstract

Parent-of-origin effect (POE) is a phenomenon whereby an allele’s effect on a phenotype depends both on its allelic identity and parent’s gender from whom the allele is inherited, as exemplified by the polar overdominance in the ovine callypyge locus and the human obesity *DLK1* locus. Systematic studies of POE of expression quantitative trait loci (eQTL) are lacking. In this study we use trios among participants in the Framingham Heart Study to examine to what extend POE exists for gene expression of whole blood using whole genome sequencing and RNA sequencing. For each gene and the SNPs in cis, we performed eQTL analysis using genotype, paternal, maternal, and joint models, where the genotype model enforces the identical effect sizes on paternal and maternal alleles, and the joint model allows them to have different effect sizes. We compared models using Bayes factors to identify paternal, maternal, and opposing eQTL, where paternal and maternal effects have opposite directions. The resultant variants are collectively called POE eQTL. The highlights of our study include: 1) There are more than 2, 000 genes harbor POE eQTL and majority POE eQTL are not in the vicinity of known imprinted genes; 2) Among 180 genes harboring opposing eQTL, 99 harbor exclusively opposing eQTL, and 58 of the 99 are phosphoprotein coding genes, reflecting significant enrichment; 3) Paternal eQTL are enriched with GWAS hits, and genes harboring paternal eQTL are enriched with drug targets. Our study demonstrates the abundance of POE in gene expression, illustrates the complexity of gene expression regulation, and provides a resource that is complementary to existing resources such as GTEx. We revisited two previous POE findings in light of our POE results. A SNP residing in *KCNQ1* that is maternally associated with diabetes is a maternal eQTL of *CDKN1C*, not *KCNQ1*. A SNP residing in *DLK1* that showed paternal polar overdominance for human obesity is a maternal eQTL of *MEG3*, offering an explanation for the baseline risk of homozygous samples through association between *MEG3* expression and obesity. Finally, we advised caution on conducting Mendelian randomization using gene expression as the exposure.

## 1 Introduction

Parent-of-origin effect (POE) is a phenomenon whereby an allele’s effect on a phenotype not only depends on its allelic identity, but also on the parent from whom the allele is inherited (DeChiara et al., 1991; Cockett et al., 1996). POE can be driven by factors such as sex bias in transmission of genetic variants (Tomé et al., 2011) and maternal genetic effects that influences the environment in which the offspring develops (Hager et al., 2008). The main driver, however, is genomic imprinting, which is a phenomenon where one of the two alleles at a locus is functionally silenced by methylation. Importantly, which parental copy of the allele being silenced is highly consistent for that gene across samples. Imprinting appears to play an important role in embryonic and placental development and social behavior (Reik and Walter, 2001; Garfield et al., 2011). Genes underpinning syndromic disorders such as Prader-Willi, Beckwtih-Weidemann, and Angelman show POE (Lawson et al., 2013). The same is also true for common disease phenotypes, such as breast cancer, diabetes, and cardiovascular disease (Kong et al., 2009; Hanson et al., 2013; Mozaffari et al., 2019).

In genetic association studies, particularly genome-wide association studies (GWAS), the absence of POE is usually assumed implicitly. This assumption is largely out of convenience, as in most GWAS, the parent-of-origin of an allele is not directly observable, and not inferable due to lack of first and second degree relatives in the sample. Exceptions include an Icelandic cohort, with their extensively documented pedigrees and the abundance of closely related samples (Kong et al., 2009), a Hutterite pedigree (Mozaffari et al., 2019), the Framingham Heart Study with three-generations of participants (Kannel et al., 1979), and the biobank datasets, for which parent-of-origin can be inferred using distant relatives (Hofmeister et al., 2022, 2025). Another contributing factor to the implicit assumption may be the presumed scarcity of documented POE in humans. Although POE is believed to be associated with a wide range of complex traits and diseases (Lawson et al., 2013), significant associations reported in GWAS setting are of limited quantity. For example, in a Hutterite study, Mozaffari et al. (2019) examined 21 phenotypes and produced 18 significant associations with POE, fewer than one association per phenotype.

Mouse studies demonstrated that, despite a limited number of imprinted genes, most phenotypes display POE, and loci that show POE were not enriched for known imprinted genes. Using reciprocal F1 crosses, Mott et al. (2014) showed that non-imprinted genes can generate POE by interactions with imprinted loci. Focusing on metabolic traits and using a mouse population at different levels of intercrossing, Macias-Velasco et al. (2022) identified a network comprised of three imprinted and six non-imprinted genes that show POE. A recent stduy on pigs demonstrated that a large number genes show POE effect in association with back fat and longissimus dorsi (Li et al., 2025). In humans, however, loci with POE in relation to phenotypes are either near imprinted regions by design (Kong et al., 2009; Hanson et al., 2013), or near regions with characteristic of imprinting from a genome-wide scan (Mozaffari et al., 2019). To what extend POE contributes to phenotypic variation in humans and how widespread the loci with POE in the human genome remains elusive.

Using whole genome sequencing and RNA sequencing data from the Framingham Heart Study, and taking advantage of its relative abundance of trios, we set to investigate to what extent and degree gene expressions is affected by parent-of-origin. There exist multiple studies on POE on gene expression (Zhabotynsky et al., 2019; Jadhav et al., 2019; Deng et al., 2020) or POE of methylation and gene expression jointly (Zink et al., 2018), but these studies are either underpowered due to small sizes or focus on analysis of allele specific gene expression instead of eQTL. A systematic study of POE eQTL is lacking and our study aims to fill this void.

## 2 Results

### 2.1 Overview of data processing

We identified 1477 trios with whole genome sequencing (WGS) data and children within the trios having RNAseq data. The trios were selected from participants in the Framingham Heart Study (FHS) who are predominantly of European descent. After routine QC of genotype data, for each trio, we first phased the child based on rules of Mendelian inheritance, which resolved all non triple heterozygous genotypes. (See Methods for details on Mendelian errors rate and how to handle them.) We then marked the remaining triple heterozygous genotypes as missing, which produced phased paternal and maternal haplotypes for each child in the trio. Next we imputed missing alleles into their paternal and maternal haplotype background, and the parental haplotype with a higher imputed allele dosage was assigned the reference allele, the other haplotype was assigned the alternative allele. Simulation studies showed this mask and impute phasing to be highly accurate (Supplementary Table S1). Thus at each SNP, we have a genotype vector, a paternal vector, and a maternal vector.

To process RNAseq data, we started from Transcript Per Million (TPM) values, and selected 16, 824 genes that have < 5% missing values (i.e., 0 TMP). We then corrected for the GC content bias using an approach that is similar to the one used by EDAseq (Risso et al., 2011). The corrected gene expression values were then quantile normalized.

### 2.2 Bayes factors as evidence for association

For genetic association we used a linear mixed model, fitted by a novel and efficient method that is designed for analyzing multiomics datasets (Guan and Levy, 2024a). The genetic relatedness matrix (twice of kinship matrix) used for the linear mixed model was estimated by Kindred (Guan and Levy, 2024c). The covariates included age, sex, body mass index, and white blood cell composition. This study focuses on genetic association between gene expression and genetic variants in cis, defined as within 1Mb from transcription start site.

We elected to use Bayes factors (Stephens and Balding, 2009) instead of p-values as the evidence of association for the following considerations. First, at many SNPs, paternal and maternal alleles have different allele frequencies, presumably due to sampling variation. P-values are not comparable unless they have the same power, while the same power requires equal allele frequencies. On the other hand Bayes factors are comparable because they take into account power through prior specifications. Second, Bayes factors panelize genetic variants with small allele frequencies to reduce false positives (Guan and Stephens, 2011; Zhou and Guan, 2018). This is useful as we observed extremely low paternal or maternal allele frequencies. Third, Bayes factors are also convenient in comparing different alternative models, such as paternal effects versus genotype effect, a setting that is difficult for p-values.

For each gene and its cis-SNP, we computed four Bayes factors: genotype Bayes factor (*BF*_*g*_), paternal Bayes factor (*BF*_1_), maternal Bayes factor (*BF*_0_), and joint Bayes factors (*BF*_*j*_). When computing joint Bayes factor, we also computed paternal and maternal effect sizes and attached a p-value to test whether they differ significantly. Examples of test statistics are provided in Table 1. *BF*_*g*_, *BF*_0_ and *BF*_1_ are one degree of freedom tests, but *BF*_*j*_ is a two degree of freedom test. Thus to achieve *BF*_*j*_ > *BF*_*g*_, the paternal and maternal effect sizes have to differ significantly to compensate for the extra degree of freedom. Examples in Table 1 are chosen because their paternal and maternal effects differ significantly, and we have *BF*_*j*_ ≫ *BF*_*g*_. Another useful observation is that *BF*_1_ is correlated with |*β*_1_|, and *BF*_0_ is correlated with |*β*_0_|. But *BF*_1_ and *BF*_0_ are oblivious to the signs of *β*_1_ and *β*_0_, and signs are informative and interesting in our context.

**Table 1:** Example of eQTL statistics. *lgBF*_*x*_ columns are log_10_ *BF*_*x*_. Column *β*_*j*_’s are effect estimates in joint analysis used to compute *BF*_*j*_. P column contains − log_10_ p-value with the null hypothesis *β*_0_ = *β*_1_ and alternative hypothesis *β*_0_ ≠ *β*_1_. Transcript start site (TSS) is in Mb, and the coordinate is from HG38. Bona fide imprinted genes are highlighted.

| Gene | TSS | SNP | $lgBF_g$ | $lgBF_1$ | $lgBF_0$ | $lgBF_j$ | $\beta_1$ | $\beta_0$ | $P$ |
| --- | --- | --- | --- | --- | --- | --- | --- | --- | --- |
| **MEG3** | 100.78 | rs12881545 | 35.7 | -0.7 | 86.3 | 85.7 | 1.0 | -21.2 | 54.0 |
| PPIEL | 39.53 | rs79473113 | 47.6 | 84.6 | 0.5 | 85.8 | -20.5 | -2.7 | 38.3 |
| **NDN** | 23.69 | rs3743340 | 36.5 | 68.5 | -0.2 | 67.7 | 18.6 | 0.8 | 33.2 |
| **PEG10** | 94.66 | rs7801134 | 2.6 | 13.3 | -0.1 | 13.6 | -8.2 | 2.4 | 12.6 |
| **ZNF331** | 53.52 | rs1284523 | 6.3 | -0.9 | 15.9 | 14.9 | -0.5 | 8.8 | 10.3 |
| ZNF890P | 5.12 | rs112844843 | 125.8 | 94.7 | 31.1 | 133.9 | 23.2 | 13.6 | 10.1 |
| **FAM50B** | 3.85 | rs111515624 | 3.5 | 11.5 | -0.5 | 10.9 | -7.4 | 0.6 | 8.6 |
| NECAB3 | 33.66 | rs2626556 | -0.7 | 1.2 | 4.6 | 5.5 | -2.9 | 4.9 | 7.8 |
| **SNURF** | 24.95 | rs4906936 | 5.1 | 12.1 | -0.9 | 11.2 | 7.7 | -0.1 | 7.6 |
| LSM7 | 2.32 | rs393651 | -1.0 | 3.5 | 1.6 | 4.9 | 4.4 | -3.3 | 7.5 |

We used log_10_ *BF* = 4 as the significance threshold. If the prior odds for a cis-eQTL is 1 out of 1000, this threshold gives the posterior probability of association (PPA) of 0.91. If the prior odds for a cis-eQTL is 1 out of 100, this threshold gives the PPA of 0.99. 1 − *PPA* is the Bayesian counterpart of the local false discovery rate (c.f. Soloff et al., 2024). We included more discussions regarding p-values, Bayes factors, and FDR in Supplementary.

### 2.3 Overview from sentinel eQTL of joint analysis

Table 1 contains 10 genes and test statistics of their sentinel eQTL. A sentinel eQTL for a gene is defined as the eQTL with largest *BF*_*j*_ for that gene. A gene with significant eQTL is called eGene. These sentinel eQTL are either *BF*_1_ ≫ *BF*_0_ ≈ 1 such as *NDN*, or *BF*_0_ ≫ *BF*_1_ ≈ 1 such as *MEG3*, or *BF*_*j*_ ≫ *BF*_*g*_ ≈ 1 such as *NECAB3*. In Table 1 the sentinel eQTL were ordered according to column *P*, which measures the significance of differences between paternal and maternal effects. Reassuringly, six out of top ten eQTL with most significant *P* are from bona fide imprinted genes (colored in blue) according to http://geneimprint.com.

Figure 1 plots paternal effect vs maternal effect for all sentinel eQTL whose log_10_ *BF*_*j*_ > 3. The color of dots corresponding to degree of difference between paternal and maternal effects. The marked sentinel eQTL include known imprinted genes such as maternally expressed *ZNF331* and *MEG3*, and paternally expressed *FAM50B, GNAS, SGCE, NDN, SNURF, SNRPN*, and *PEG10*. Imprinted genes are located along the x and y-axes. At least two genes appear to be novel imprinted genes: *PPIEL* and *CCR9*. Gene *ZNF890P* provides an example of paternal and maternal effects being in the same direction, but their sizes differ significantly. Finally, *NECAB3* and *LSM7* are two examples of paternal and maternal effects that are in opposite directions.

**Figure 1:**
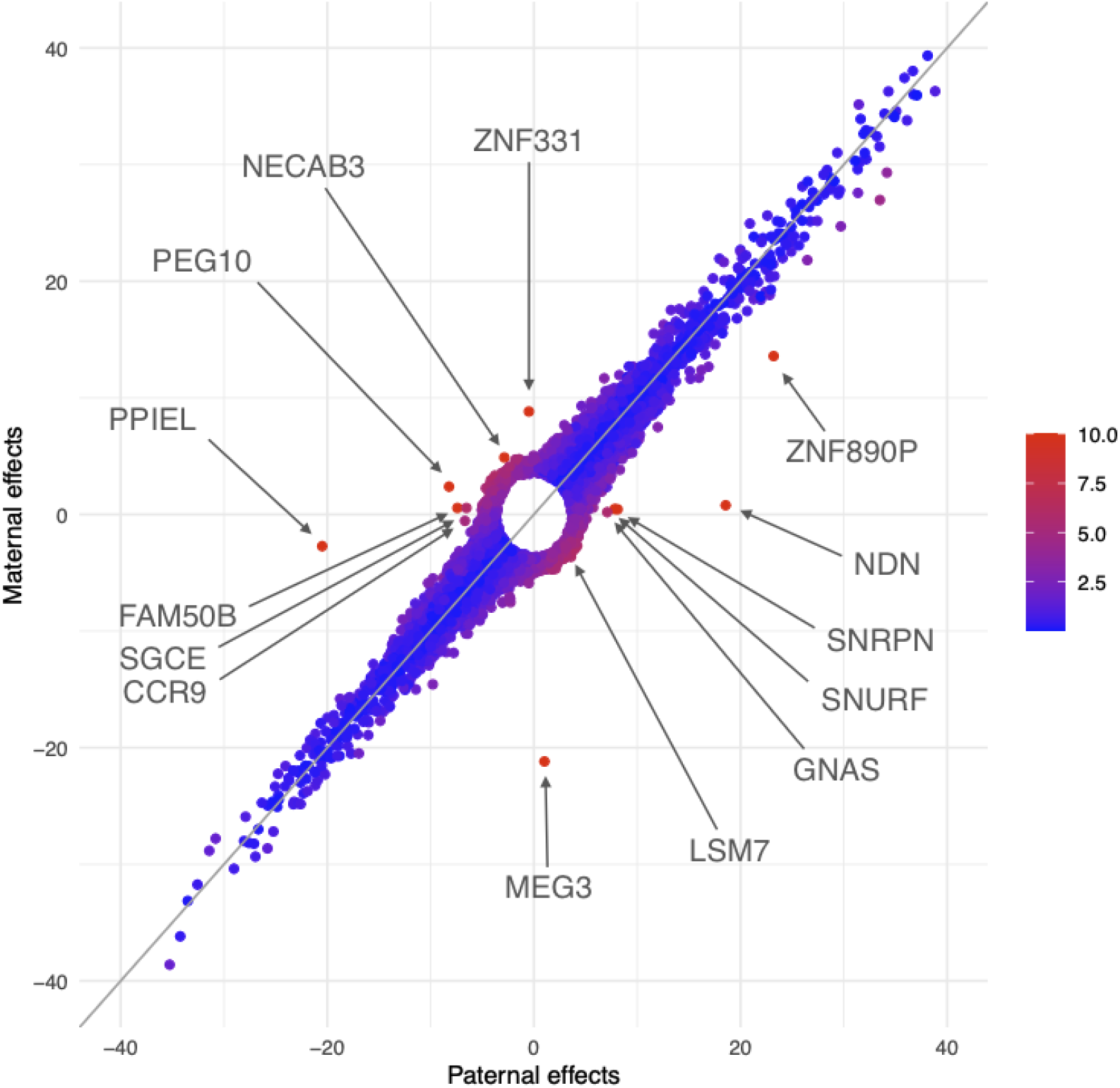
Comparison of paternal effects and maternal effects of sentinel eQTL in joint analysis. Each point is the most prominent eQTL with log_10_ *BF*_*j*_ > 3 of gene. The paternal effect is on x-axis and maternal effect y-axis. The coloring reflects significance of differentials between paternal effects and maternal effects, with red more significant than blue.

This approach based on the sentinel eQTL suggests that 1) the combined effect of paternal and maternal allele can be across all 360 degrees; 2) when paternal and maternal effects are in opposite directions, both effects sizes are modest compared to when they are in the same direction; and 3) there are quite a few sentinel eQTL whose paternal and maternal effects show opposite directions.

### 2.4 *NECAB3* : opposite paternal and maternal effects

The existence of eGenes whose eQTL show opposite paternal and maternal effects is intriguing. We focus on the example of *NECAB3* to examine further details. Figure 2 left panel shows that all eQTL of *NECAB3* have opposite paternal and maternal effects. Consequently, the joint Bayes factors *BF*_*j*_’s are much larger than their corresponding genotype Bayes factors *BF*_*g*_’s (right bottom). We select as an example SNP *rs4911348*, which is different from the sentinel eQTL shown in Figure 1, and looked into details of its association with the gene expression of *NECAB3* (phenotype). Three panels of boxplots (right top) show that genotype has no association with the phenotype, and both paternal alleles and maternal alleles are associated with the phenotype, but in opposite directions. Opposite paternal and maternal effects were also observed in Hutterite POE study, where ten associations of opposite effects across nine traits were reported (Mozaffari et al., 2019).

**Figure 2:**
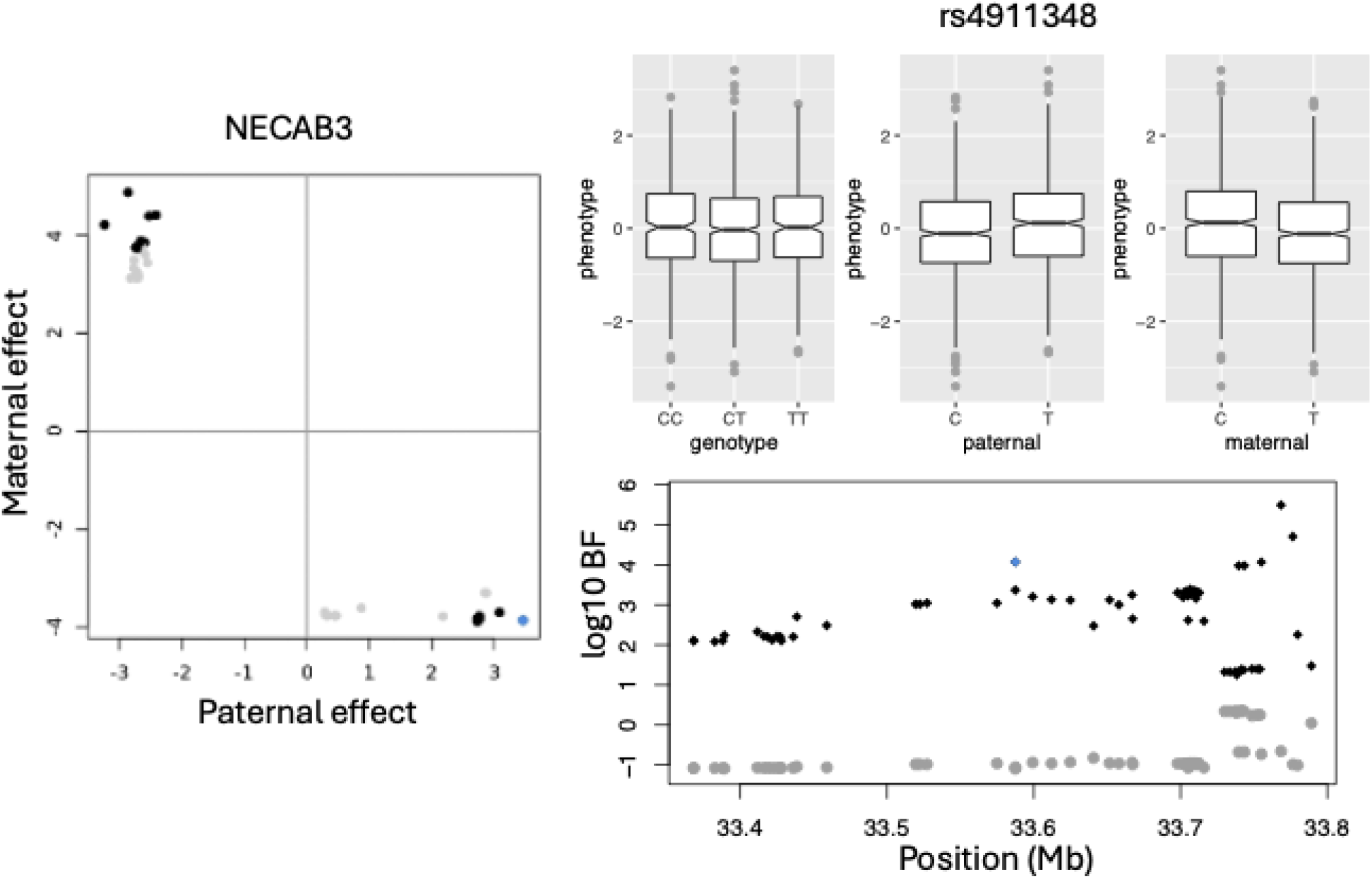
An example of gene that whose paternal and maternal eQTL have opposite effect sizes. On the left is paternal vs maternal effect sizes normalized by their corresponding standard deviation. The effect sizes were estimated by joint analysis. The non-significant eQTL are colored in gray and significant eQTL colored in black. An eQTL marked in blue was chosen to show boxplots with gene expression (phenotype) of NECAB3. The three panels of boxplots are for genotypes, paternal alleles, and maternal alleles. For this SNP rs4911348, the genotypes has no association with phenotype, but both paternal alleles and maternal alleles are associated with the phenotype, and the paternal and maternal effects are in opposite direction. Right panel bottom show Manhattan plot of cis-eQTL of the gene NECAB3. The genotype Bayes factor were colored in gray and joint test Bayes factor were colored in black. SNP rs4911348 was highlighted in blue.

### 2.5 POE eQTL are abundant

Next we looked beyond the sentinel eQTL and examined all significant eQTL. We noted that there are 15, 893 eQTL from 14, 733 SNPs and 1, 824 eGenes that are insignificant for the SNP test, but significant for the paternal, or maternal, or joint test. That is, log_10_ *BF*_*g*_ < 4, but either log_10_ *BF*_1_ > 4, or log_10_ *BF*_0_ > 4, or log_10_ *BF*_*j*_ > 4. In other words, these eQTL cannot be detected without analyzing POE, which implies that POE can contribute to recover missing heritability.

The minor allele frequencies of the POE eQTL are balanced. Supplementary Figure S4 empirically demonstrates that our findings are not disproprtionately driven by low-frequency variants and provide a more complete picture of the genetic architecure underlying these POE signals.

With reference Figure 1, we proposed criteria to define subsets of eQTL. The first subset eQTL locate along the x-axis, mimicking sentinel eQTL of *NDN* and *FAM50B* ; the second subset locate along the y-axis, mimicking sentinel eQTL of *ZNF331* and *MEG3* ; the third subset locate along the secondary diagonal, mimicking sentinel eQTL of *NECAB3* and *LSM7* ; and the fourth subset follows (blues dots) along the diagonal line. Note Figure 1 only involves *BF*_*j*_, our criteria to define gene sets also takes into account of *BF*_1_, *BF*_0_, and *BF*_*g*_ (details in Methods).

The first set of eQTL have significant paternal effects but insignificant maternal effects. We identified 15, 576 such *paternal eQTL* (set *S*_*P*_) from 14, 372 SNPs associated with 1, 188 eGenes. The second set of eQTL have significant maternal effects but insignificant paternal effects. We identified 14, 783 such *maternal eQTL* (set *S*_*M*_) from 13, 293 SNPs associated with 1, 209 eGenes. We refer to these paternal and maternal eQTL as *imprinting eQTL*. The third set of eQTL have opposite paternal and maternal effects, such that joint Bayes factors are much larger than genotype Bayes factor *BF*_*j*_ ≫ *BF*_*g*_. We identified 688 such *opposing eQTL* (set *S*_*O*_) from 485 SNPs that are associated with 180 eGenes. Imprinting eQTL and opposing eQTL are referred to as *POE eQTL*.

The fourth set of eQTL require that both paternal and maternal alleles alone are associated with an expression phenotype, and the two effects are in the same direction. For these eQTL *BF*_*g*_ > *BF*_*j*_ because similar effect sizes favor one degree of freedom test. We identified 884, 119 such *genotype eQTL* from 577, 701 SNPs and 4940 eGenes. The percent of eGenes with genotype eQTL is on par, but smaller than GTEx study (Consortium et al., 2020), presumably because log_10_ *BF*_*g*_ > 4 is a more stringent threshold for eQTL analysis.

### 2.6 Paternal eQTL are enriched with GWAS hits

Using ANNOVAR (Wang et al., 2010), we annotated all four sets of eQTL (Table 2). We made the following observations: 1) The median distance to transcription start site (TSS) for opposing eQTL is 673.2Kb, much larger than other types of eQTL, and majority of opposing eQTL *S*_*O*_ are located in introns of nearby genes. 2) The percent of GWAS hits for SNPs in *S*_*O*_ is 0.022, significantly lower than that of SNPs in genotype eQTL *S*_*G*_’s 0.054 (test for proportion *P* = 0.001). This is reasonable as GWAS mainly use genotype test and are agnostic to opposing eQTL. 3) The precent of GWAS hits for SNPs in paternal eQTL *S*_*P*_ is higher than that of SNPs in *S*_*G*_ (test for proportion *P* = 9 *×* 10^−11^), and the percent of GWAS hits among SNPs in maternal eQTL *S*_*M*_ is similar to that of SNPs in *S*_*G*_ (test for proportion *P* = 0.051).

**Table 2:** Top tabular: Annotation of eQTL. *S*_*O*_ is a set of opposing eQTL, *S*_*P*_ is a set of paternal eQTL, *S*_*M*_ is a set of maternal eQTL, and *S*_*G*_ is a set of genotype eQTL. The column eQTL contain counts of eQTL in each set, and column SNP contains counts of distinct SNPs of eQTL in the set. The column *eGene* is the number of genes associated with eQTL in the set. The column of Len is the median length in Kb of the eGenes. (The pattern is the same with the mean length.) The column *D*_*TSS*_ contains median distance in Kb to transcription start site. The column GWAS contains percent of GWAS hits among SNPs, with GWAS p-value threshold of 5 *×* 10^−8^. Bottom tabular: A SNP set *P*_*k*_ contain SNPs in *S*_*G*_ that are eQTL of at least *k* eGenes. (Note *P*_1_ = *S*_*G*_.) Percent of GWAS hits for SNP set *P*_*k*_ increases with *k*.

| Set | eQTL | eGene | Len | $D_{TSS}$ | Genic | Intron | Exon | ncRNA | SNP | GWAS |
| --- | --- | --- | --- | --- | --- | --- | --- | --- | --- | --- |
| $S_O$ | 688 | 180 | 28.1 | **673.2** | 0.260 | **0.663** | 0.006 | 0.026 | 485 | **0.023** |
| $S_P$ | 15,576 | 1188 | 23.3 | 122.9 | 0.362 | 0.490 | 0.016 | 0.080 | 14,372 | **0.066** |
| $S_M$ | 14,783 | 1209 | 23.8 | 122.5 | 0.365 | 0.487 | 0.016 | 0.076 | 13,293 | 0.050 |
| $S_G$ | 884,119 | 4940 | 24.6 | 84.1 | 0.383 | 0.449 | 0.017 | 0.091 | 577,701 | 0.054 |
| $P_1$ | 884,119 | 4940 | 24.6 | 84.1 | 0.383 | 0.449 | 0.017 | 0.091 | 577,701 | 0.054 |
| $P_2$ | 471,354 | 2855 | 17.7 | 96.8 | 0.400 | 0.403 | 0.019 | 0.112 | 164,936 | 0.071 |
| $P_3$ | 269,546 | 1536 | 13.8 | 114.4 | 0.440 | 0.349 | 0.020 | 0.129 | 64,032 | 0.088 |
| $P_4$ | 172,871 | 858 | 11.0 | 134.2 | 0.488 | 0.295 | 0.017 | 0.143 | 31,807 | 0.097 |
| $P_5$ | 126,107 | 510 | 8.6 | 143.8 | 0.508 | 0.284 | 0.016 | 0.140 | 20,116 | 0.110 |
| $P_6$ | 94,367 | 383 | 7.7 | 146.7 | 0.520 | 0.284 | 0.015 | 0.135 | 13,768 | 0.117 |
| $P_7$ | 46,925 | 272 | 6.1 | 131.8 | 0.593 | 0.234 | 0.015 | 0.106 | 5,861 | 0.133 |
| $P_8$ | 26,695 | 178 | 3.7 | 116.6 | 0.619 | 0.221 | 0.010 | 0.099 | 2,971 | 0.144 |
| $P_9$ | 12,783 | 124 | 2.3 | 121.8 | 0.483 | 0.355 | 0.011 | 0.111 | 1,232 | 0.198 |

Number of eQTL per SNP (*r*) measures degree of pleiotropy of a set of SNPs. SNPs in *S*_*P*_ and SNPs in *S*_*M*_ have similar pleiotropy *r* ≈ 1.1, SNPs in *S*_*G*_ have the largest *r* = 1.53, and SNPs in *S*_*O*_ display *r* = 1.41. In other words, SNPs are more exclusive for imprinting eQTL, less exclusive for genotype eQTL, and somewhere in between for opposing eQTL. Table 2 (bottom) compares how the degree of pleiotropy affects their annotation for SNPs in *S*_*G*_, where set *P*_*k*_ contains SNPs in *S*_*G*_ that are eQTL of at least *k* eGenes. The most interesting observation is that the percent of GWAS hits increases with rising SNP pleiotropy. This feature is only observed in genotype eQTL.

### 2.7 POE eGenes are abundant

There are 1, 188 eGenes harboring paternal eQTL and 1, 209 eGenes harboring maternal eQTL. Their union contains 2, 139 eGenes harbor imprinting eQTL, and their intersection contains 258 eGenes harboring both paternal and maternal eQTL. There are 4, 940 eGenes harboring genotype eQTL, and 1, 867 eGenes harboring both genotype eQTL and imprinting eQTL. Supplementary Figure S1 showed an example of an eGene harboring both maternal and paternal eQTL: *GZMH*, and an example of an eGene harboring both genotype eQTL and imprinting eQTL: *DSE*. The most prominent feature is that paternal eQTL and maternal eQTL cluster in separate genomic regions, and genotype eQTL and imprinting eQTL also cluster in separate genomic region.

### 2.8 eGenes that harbor exclusively imprinting eQTL

There are 129 eGenes (set *G*_1_) that harbor exclusively paternal eQTL, and 139 eGenes (set *G*_0_) that harbor exclusively maternal eQTL. Naturally, *G*_1_ contain many bona fide imprinted genes with paternal expression such as *FAM50B, NDN, SGCE, SNRPN, SNURF* and *PEG10*. Remaining genes in *G*_1_ are candidates for imprinted genes with paternal expression, for example *CCR9* and *NCOA2. G*_0_ contain bona fide imprinted genes with maternal expression such as *GRB10, ZNF331* and *MEG3*. The remaining in *G*_0_ are candidates for imprinted genes with maternal expression, for example *COA8* and *ZNF888*.

The Supplementary Material provides a full list of the candidate imprinted genes from the eQTL analysis. Note however, that not all imprinted genes have eQTL, therefore this list is incomplete. Moreover, a gene on the list is considered behaving like an imprinted gene, but may not be a bona fide imprinted gene. For example, a gene that has no cis-eQTL on its own, may “acquire” a cis-eQTL through gene-gene interaction with a putative imprinted gene such that the cis-eQTL is in fact its trans-eQTL.

We used *G*_1_ and *G*_0_ as proxies for paternal and maternal imprinted genes, and used cytoBands these genes occupies as proxies for imprinted regions to examine to what extend paternal and maternal eQTL locate in the same cytoBands as these proxy imprinted genes. There are 811 autosome cytoBands in humans, 765 of them larger than 1 Mb, and the median size is 3.2 Mb (Cheung et al., 2001). The 129 proxy paternal imprinted genes occupy 89 cytoBands (set 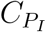), while 15, 576 paternal eQTL occupy 445 cytoBands (set 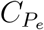), among them 9, 947 (or 64%) were not in the cytoBands set 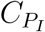. On the other hand, the 139 proxy maternal imprinted genes occupy 96 cytoBands (set 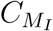), and 14, 783 maternal eQTL occupy 465 cytoBands (set 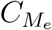), among them 7, 987 (or 54%) were not in the cytoBand set 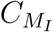.

Therefore, a majority of imprinting eQTL are not in the vicinity of the proxy imprinted regions. Interestingly, 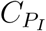 and 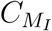 share only 14 cytoBands, 8% of their union of 171 cytoBands. As a comparison, 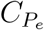 and 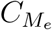 share 349 cytoBands, 62% of their union of 561 cytoBands.

### 2.9 eGenes that harbor exclusively opposing eQTL

There are 99 eGenes (set *G*_2_) that harbor only opposing eQTL (Supplementary). Among them 58 encode phosphoproteins (Supplementary), a significant enrichment (FDR = 5.5 *×* 10^−4^) according to DAVID, a web server for functional enrichment analysis (Sherman et al., 2022). A phosphoprotein is a protein that is posttranslationally modified by the attachment of either a single phosphate group, or a complex molecule such as 5’-phospho-DNA, through a phosphate group. Because phosphorylation often serves as an on/off switch, targeting phosphoproteins or the enzymes regulating them can restore normal cell signaling in diseases. In addition, drugs can modulate the functions of these proteins, either inhibiting or enhancing their activities. This makes phosphoprotein attractive drug targets. Indeed, according to a therapeutic target database (Zhou et al., 2024), among 58 phosphoproteins genes that harbor exclusively opposing eQTL, there are three successful drug targets *ITPR1, ITPR2*, and *REL*, and three additional clinical trial targets *CD46, NOTCH3*, and *XPO1*.

Other notable phosphoproteins genes include *CDKN2A, PHIP*, and *LEP*, among others. Where *CDKN2A* produces two major proteins p16(*INK4*), which is a cyclin-depnednet kinase inhibitor, and p14(*ARF*), which binds the p53-stabilizing protein MDM2 (Serrano et al., 2000); *PHIP* is associated with Chung-Jansen syndrome, featuring behavioral problems, intellectual disability, obesity, and dysmorphia (Webster et al., 2016; Jansen et al., 2018); and *LEP* encodes leptin, a protein that plays a critical role in the regulation of body weight. Leptin is secreted by white adipocytes, and it binds to the leptin receptor in the brain, which in turn inhibits appetite and promotes energy expenditure (Maffei et al., 1995; Weigle et al., 1997).

### 2.10 Paternal eGenes are enriched with drug targets

We used a therapeutic target database (Zhou et al., 2024) to examine the enrichment with drug targets of the four sets of eGenes, grouped by their eQTL. Table 3 contains the enrichment of the four gene sets in either successful targets, clinical trial targets, or combined targets. The significant enrichment that (following Bonferroni correction of 12 tests) is highlighted. Both paternal eGenes and genotype eGenes are enriched in clinical trial targets and combined targets, but paternal eGenes are significantly more enriched than genotype eGenes (one-sided test for proportion *P* = 0.035). The enrichment of drug targets for paternal eGenes echoes the enrichment of GWAS hits for paternal eQTL (Table 2). This asymmetry between paternal and maternal eGenes perhaps finds its root in the conflicting interests of two sets of genes in relation to transfer of nutrients from the mother to her offspring (Moore and Haig, 1991; Hitchcock and Gardner, 2019). In maze, gene expression of the hybrid is regulated exclusively by the paternally transmitted alleles (Swanson-Wagner et al., 2009).

**Table 3:**
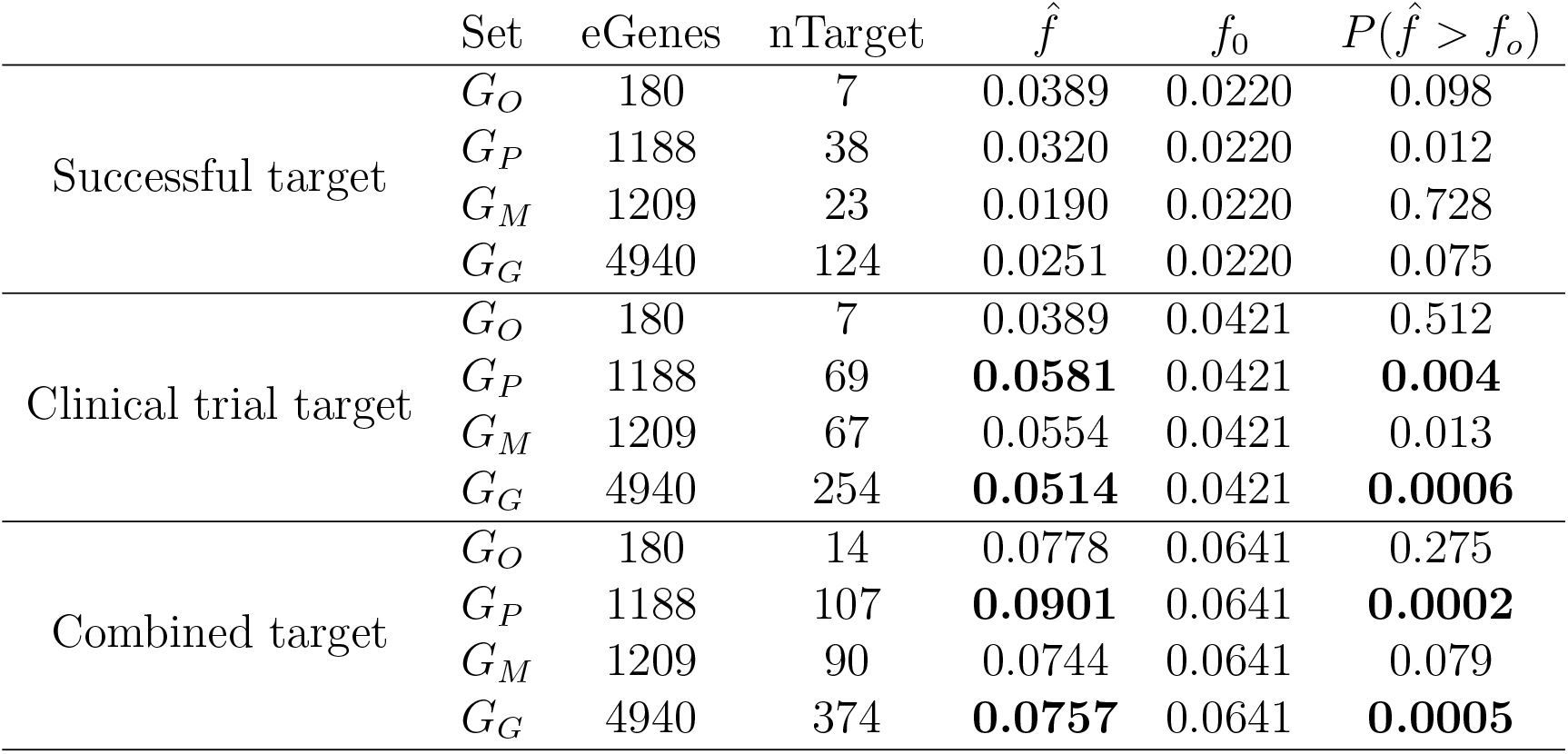
Enrichment of drug target genes. *G*_*O*_ contains eQTL with both paternal and maternal effects but they are in opposite direction. *G*_*P*_ contains eQTL of paternal effects. *G*_*M*_ contains eQTL of maternal effects. *G*_*G*_ contain eQTL that have both paternal and maternal effects and they are in the same direction. Column nGene are sizes of gene sets, Column nTarget are counts of successful drug targets. The last column contains one-sided test-of-proportion p-values. The highlighted p-values are significant after Bonferroni correction of 12 tests.

| | Set | eGenes | nTarget | $\hat{f}$ | $f_0$ | $P(\hat{f} > f_0)$ |
| --- | --- | --- | --- | --- | --- | --- |
| Successful target | $G_O$ | 180 | 7 | 0.0389 | 0.0220 | 0.098 |
| | $G_P$ | 1188 | 38 | 0.0320 | 0.0220 | 0.012 |
| | $G_M$ | 1209 | 23 | 0.0190 | 0.0220 | 0.728 |
| | $G_G$ | 4940 | 124 | 0.0251 | 0.0220 | 0.075 |
| Clinical trial target | $G_O$ | 180 | 7 | 0.0389 | 0.0421 | 0.512 |
| | $G_P$ | 1188 | 69 | **0.0581** | 0.0421 | **0.004** |
| | $G_M$ | 1209 | 67 | 0.0554 | 0.0421 | 0.013 |
| | $G_G$ | 4940 | 254 | **0.0514** | 0.0421 | **0.0006** |
| Combined target | $G_O$ | 180 | 14 | 0.0778 | 0.0641 | 0.275 |
| | $G_P$ | 1188 | 107 | **0.0901** | 0.0641 | **0.0002** |
| | $G_M$ | 1209 | 90 | 0.0744 | 0.0641 | 0.079 |
| | $G_G$ | 4940 | 374 | **0.0757** | 0.0641 | **0.0005** |

## 3 Discussion

We performed parent-of-origin analysis of eQTL in whole blood gene expression using 1477 trios from the Framingham Heart Study. Some highlights of our study include: 1) There are more than 2,000 genes harboring POE eQTL and majority POE eQTL are not in the vicinity of known imprinted genes; 2) Among 180 genes harboring opposing eQTL, 99 harbor exclusively opposing eQTL, and 58 of the 99 are phosphoprotein coding genes, reflecting significant enrichment; 3) Paternal eQTL are enriched with GWAS hits, and genes harboring paternal eQTL are enriched with drug targets.

Although our study lacks replications in independent cohorts, we have taken several steps to ensure the internal validity and robustness of our findings within the current study. Specifically, we used well-established, *bona fide* imprinted genes (e.g., *MEG3, NDN, SNURF*, as listed in Table 1 and Figure 1) as positive control. We used conservative calling creiteria, and explored different parameters (Supplementary Table S2) and chose the one that minimized the inclusion of false positives, ensuring that our core gene sets (e.g., *G*_1_, *G*_0_) are high-confidence discoveries. We developed a rigorous analytical pipeline, including traio-based phasing validated by simulation (Supplmentary Table S1), the use of linear mixed models to control for relatedness and population structure, and the application of Bayes factors which inheritenly panalize variants with low minor allele frequencies, thereby reducing spurious associations. We believe these internal consistency checks and methodological rigor provide strong confidence in our findings.

Our study demonstrated abundance of POE in gene expression, illustrated additional complexity of gene expression regulation, and provided a resource that is complementary to existing resources such as GTEx. We conclude by revisiting two previous findings in light of our work, and advising caution on conducting Mendelian randomization using gene expression as the exposure.

### 3.1 T2D and maternal allele of rs2237892

SNP rs2237892 in the last intron of *KCNQ1* was found in association with type 2 diabetes (T2D) in a Japanese cohort (Yasuda et al., 2008), and the association was later confirmed in a Chinese cohort (Liu et al., 2009). Three other SNPs (rs2283228, rs2237895, and rs2237897) in the same last intron of *KCNQ1* were found in association with T2D in another Japanese cohort (Unoki et al., 2008). Both rs2237895 and rs2237897 were replicated in a Singaporean cohort, a Danish cohort (Unoki et al., 2008), and a Chinese cohort (Liu et al., 2009). In these studies, *KCNQ1* is identified as T2D candidate gene.

Based on these results from the genotype test, Kong et al. (2009) demonstrate that SNP rs2237892 is maternally associated with T2D in an Iceland cohort. This maternal association was replicated with a larger odds ratio in a Pima Amerindian cohort (Hanson et al., 2013). According to data from GTEx, none of these SNPs are eQTL of *KCNQ1* and *CDKN1C* in any tissue, consistent with the fact that these are imprinted genes and GTEx study is oblivious to parent of origin. Our eQTL analysis showed that SNPs rs2237892, rs2283228, and rs2237897 are maternal eQTL of *CDKN1C*, a down-stream neighbor of *KCNQ1*, and none of these SNPs are eQTL of *KCNQ1*.

We therefore suggest that *CDKN1C*, instead of *KCNQ1*, is a T2D candidate gene, for the following reasons: 1) These GWAS SNPs connect T2D and *CDKN1C* quantitatively, but connect T2D and *KCNQ1* only geographically; 2) A study suggests that *CDKN1C* mutations may represent a novel monogenic form of diabetes (Kerns et al., 2014); 3) A boy carrying a frameshift mutation in *CDKN1C* was diabetic from week 29 (Berland et al., 2022); and 4) Targeted demethylation at the CDKN1C/p57 locus induces human *β* cell replication, while the loss of insulin-secreting *β* cell is characteristic among T1D and T2D Ou et al. (2019).

We note that gene expression in our study is from whole blood. While whole blood is a valuable and accessible tissue, replication in T2D-relevant tissues (e.g., pancreas, adipose) would be an important future direction, and our findings provide a hypothesis for such targeted investigations.

### 3.2 Polar overdominance and rs1802710

Heterozygous overdominance is a pattern of inheritance such that both homozygous AA and BB have the same baseline trait value, while heterozygous AB has a higher trait value. Polar overdominance (Cockett et al., 1996) introduces asymmetry into overdominance to separates AB into two groups according to parent-of-origin of A (or B), such that AB with paternal A (or B) has a high trait value, while AB with maternal A (or B) has the baseline trait value.

Wermter et al. (2008) studied trios of extremely obese offspring and identified rs1802710 in exon 5 of *DLK1*, homologous to the ovine callipyge locus (Cockett et al., 1996), whose allelic transmission pattern was consistent with polar overdominance: frequent transmission of the paternal C allele to obese children, but the relative risk for carriers of the homozygous CC genotype was not increased compared to the reference TT genotype.

Our eQTL results show that *DLK1* has no significant eQTL, but the maternal C allele of rs1802710 reduces expression of *MEG3*, 53Kb downstream of *DLK1*. Both *MEG3* and *DLK1* are imprinted with *MEG3* maternally expressed and *DLK1* paternally expressed. The expression of *MEG3* was shown to be significantly higher in the obese group (Daneshmoghadam et al., 2021). Therefore the maternal C allele is associated with reduced risks of obesity, which balances out the increased risk conferred by paternal C allele, thus offering an explanation for the baseline risk of homozygous CC samples in (Wermter et al., 2008). Since rs1802710 is not an eQTL of any gene in any tissue according to GTEx data, our study is critical to link rs1802710 with *MEG3*, albeit not in the adipose tissue.

### 3.3 Mendelian randomization

Mendelian randomization (MR) refers to the random allocation of alleles at the time of gamete formation. Observational epidemiology studies use MR to infer the causal effect of an exposure on an phenotype (Smith and Ebrahim, 2003), as if the random allocation of alleles is comparable to a randomized clinical trial. An important assumption for MR, among many other important assumptions, is that the phenotype conferred by a specific genetic variant is homogeneous in the population, and exchangeable between paternally and maternally inherited alleles. POE is a direct violation of this assumption (Bochud et al., 2008).

There is a growing interest in using gene expression as an exposure to perform MR across various traits (Porcu et al., 2019; van der Graaf et al., 2020). If an exposure has eQTL with POE, ignoring the parent-of-origin will bias prediction of a subset of samples. If the bias is severe it will change the ranking of the exposure. Consequently, it either compromises the power of the MR or leads to a false conclusion of a causal relationship between the exposure and the phenotype.

Our study demonstrates the abundance of POE in gene expressions, and we therefore advise caution when conducting MR using gene expression as the exposure. We suggest checking the list of POE SNPs and eGenes we provided in the Supplementary Material and exclude those that show POE towards the exposure. The same suggestion is also applicable to computing polygenic risk scores, and imputing gene expression such as PrediXcan (Gamazon et al., 2015) and a Bayesian method motivated by it (Qi et al., 2018).

## 4 Method

### 4.1 Genotype data

We first used pedigree and availability of RNAseq data to narrow the samples down to 2955, and performed routine QC to obtain bi-allelic SNPs (Taliun et al., 2021). We then used Kindred to infer pairwise kinship for each chromosome, and computed mean and sd of the kinship. A pair of sample that is parent-offspring can be distinguished from full sibs by the sd (Supplementary Figure S2). Trios are consisted of sample A, B, and C such that *ϕ*(*A, B*) ≈ 0 and *ϕ*(*A, C*) ≈ *ϕ*(*B, C*) ≈ 0.25 where *ϕ*(*·, ·*) is the kinship. In the end, we identified 1477 trios that have whole genome sequencing data and whose children have RNAseq data. Totally, we have 7, 752, 281 autosome bi-allelic SNPs with minor allele frequency > 0.01 (not corrected for relatedness). Across all 1477 trios, the Mendelian error is negligible 4.76 per 100, 000 SNPs per trio. To handle Mendelian errors, we assume child was correctly genotyped, and errors happened in parents, but if the child is heterozygous, we assume this is a triple heterozygous SNP. On average 1477 trios have 65 triple heterozygous genotypes per SNP. The triple heterozygous SNPs cannot be phased by Mendelian inheritance, but can be phased based on a linkage disequilibrium model (below).

### 4.2 Inference of parent of origin

For each trio, we first phase all markers that are not triple heterozygous using rule of Mendelian inheritance, where for the child in each trio we obtain paternal and maternal haplotypes with triple heterozygous markers whose phase are unresolved. We mark those triple heterozygous markers as missing. For markers that show Mendelian incompatibility, we assume child’s genotype is correct and mark as missing if it is heterozygous. Using a hidden Markov model that was designed to model haplotype variation (Guan, 2014), we impute missing markers in each haplotypes to obtain dosage estimates (between 0 and 1). Then for each marker that are marked as missing, we compare imputed dosages between paternal and maternal haplotypes, and assign the large dosage as 1 and small dosage as 0. We thus obtain paternal and maternal haplotypes with no missing data. We simulated trios using haplotypes from 1000 Genomes project, and investigated the accuracy of the phasing by mask and imputation approach described above. The phasing is highly accurate, the error rate is less than 1 out of 200 triple heterozygous SNPs (Supplementary Table S1).

### 4.3 RNAseq data

The RNAseq data of 1477 chilren comes from three batches with 1381 comes from batch 1; 14 comes from batch 2, and 82 comes from batch 3. Our primary goal is to correct for GC content bias. To this end, we first remove genes whose proportion of 0 TPM values is greater than 5%, which retain 16, 969 genes out of total 58, 103 transcripts, majority of which are coding genes. Among those, 16, 824 genes have GC content. To correct for GC content bias, we took an approach used by EDAseq to regress out the GC content from log(1+*TMP*), but instead of using local linear regression as documented in EDAseq, we used local quadratic regression implemented in loess in R. Specifically, let *y* = log(1+*TMP*) be a gene expression and *g* be the GC content, we fit *ly* = *loess*(*y, g*) and compute *ŷ* = *median*(*y*) + *ly*$*residuals*. We then quantile normalize *ŷ* by *qqnorm*(*ŷ, plot*.*it* = *F*)$*y* separately for each gene.

### 4.4 eQTL analysis

We used IDUL to perform eQTL analysis. IDUL fits linear mixed models to achieve exact optimal. It was specifically developed to analyze multi-omics data to achieve high efficiency by reusing the intermediate computations (Guan and Levy, 2024a). Linear mixed model includes a random effect with a covariance structure defined by the genetic relatedness matrix (GRM). This approach is widely regarded as more robust than including a limited number of principal components, as it accounts for both fine-scale population stratification and known relatedness simultaneously. Previous analysis showed that for Framingham heart study, the genomic inflation was well controlled using linear mixed model with kinship matrix computed by Kindred (Guan and Levy, 2024c). In this study, we also controlled age, sex, BMI, and cell compositions in peripheral blood. We now briefly describe how Bayes factors were computed.

Consider a model

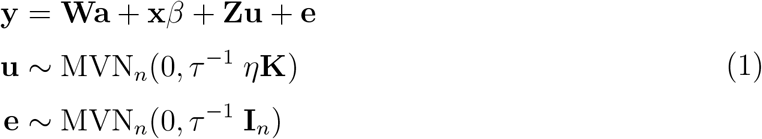

where **W** contains conventional covariates such as age and sex, including a column of 1, **x** contains genetic variant(s) to be tested for association, **u** is the random effect with **Z** as its loading matrix and twice of kinship **K** as its covariance (both **Z** and **K** are known), MVN_*n*_ denotes an n-dimensional multivariate normal distribution, **I**_*n*_ is n-dimensional identity matrix. In our anlaysis **Z** is the identity matrix. Denote **X** = (**W, x**) and **b** = (**a**, *β*), then **Xb** is the fixed effect, and we assume **X** has a full rank *c*. In genetic association studies, the random effect **Zu** is a nuisance term that absorbs part of the phenotype **y** that is attributable to population stratification and relatedness. The maximum likelihood estimate (MLE) of *η* can be efficiently obtained (Guan and Levy, 2024a), plug 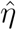 back into (1), and specify the following conjugate prior

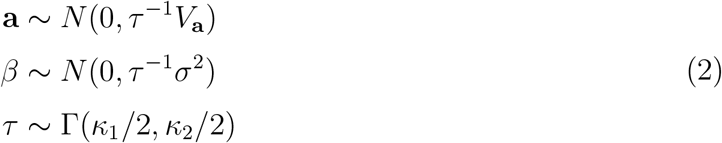

and let *V*_**a**_ → ∞, *κ*_1_ → 0 and *κ*_2_ → 0, Bayes factor can be evaluated efficiently in a closed form. This study we used *σ* = 0.5, following prior work of Bayes factors for linear models (Servin and Stephens, 2007). Complete details on computing Bayes factors for linear mixed model can be found in (Guan and Levy, 2024b).

### 4.5 P-value for difference between paternal maternal effects

For joint analysis, we have

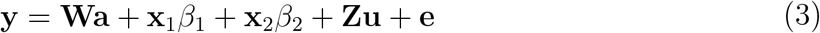

with the prior for *β*_1_ and *β*_2_ as

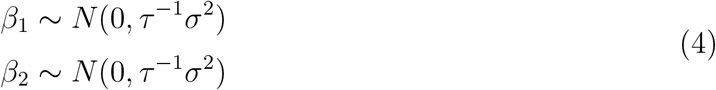

The Bayes factor can be similarly computed in a closed form. We obtain the posterior estimates of 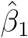 and 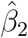, their variances 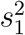 and 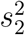 and covariance *s*_12_. To test null hypothesis 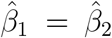, we compute test statistics 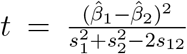. Since under the null *t* following *χ*^2^ distribution with 1 degree of freedom, we can compute a p-value.

### 4.6 Threshold for eQTL sets and gene sets

For paternal eQTL set *S*_*P*_, we require log_10_ *BF*_1_ > 4 and log_10_ *BF*_0_ < *θ*; For maternal eQTL set *S*_*M*_, we require log_10_ *BF*_0_ > 4 and log_10_ *BF*_1_ < *θ*; For opposing eQTL set *S*_*O*_, we require log_10_ *BF*_*j*_ − log_10_ *BF*_*g*_ > 4 and log_10_ *BF*_1_ > *θ* and log_10_ *BF*_0_ > *θ*, and *β*_1_ ∗ *β*_0_ < 0; For genotype eQTL set *S*_*G*_, we require log_10_ *BF*_*g*_ > 4 and log_10_ *BF*_1_ > *θ* and log_10_ *BF*_0_ > *θ*, and *β*_1_ ∗ *β*_0_ > 0. It’s easy to see for a particular type of eQTL, different *θ* produce nested eQTL sets. We tried *θ* = 0, log_10_ 2, and log_10_ 3 to obtain eQTL sets (Supplementary Table S2), from which we obtain gene sets harboring those eQTL *G*_*P*_, *G*_*M*_, *G*_*A*_ and *G*_*G*_, from which we obtain genes harbor exclusively paternal eQTL *G*_1_ = *G*_*P*_ \ (*G*_*M*_ ∪ *G*_*A*_ ∪ *G*_*G*_), genes harbor exclusive maternal eQTL *G*_0_ = *G*_*M*_ \ (*G*_*P*_ ∪ *G*_*A*_ ∪ *G*_*G*_), and genes harbor exclusively opposing eQTL *G*_0_ = *G*_*A*_ \ (*G*_*P*_ ∪ *G*_*M*_ ∪ *G*_*G*_). We compare known imprinted genes with paternal expression with *G*_1_, and known imprinted genes with maternal expression with *G*_0_. *θ* = log_10_ 2 produced minimum *G*_1_ and *G*_0_ that contain known imprinted genes.

## 1 Simulation study of phasing trios

There are 503 European (EUR), 504 East Asian, and 661 African (AFR) phased samples in 1000 Genomes project. Our simulation was conducted separately for each population. In each population, we randomly selected two samples without replacement and assign the first as father and the second as mother, until exhausted all samples or had only one sample left. Using Chromosome 22, we selected one haplotype from the father and one from the mother to simulate the child. We didn’t simulate genotyping error, as in real data the Mendelian error is negligibly small. We then ran the process of phasing by rule of Mendelian inheritance for each trio, masked triple heterozygous SNPs for each child, jointly fit an LD model using all chidren’s haplotypes to impute masked SNPs, and finally based on imputed allele dosage to assign alleles (reference or alternative) to maternal and paternal haplotypes. We tallied all counts and put in Table S1, and results show our approach of phasing is highly accurate.

**Table S1:**
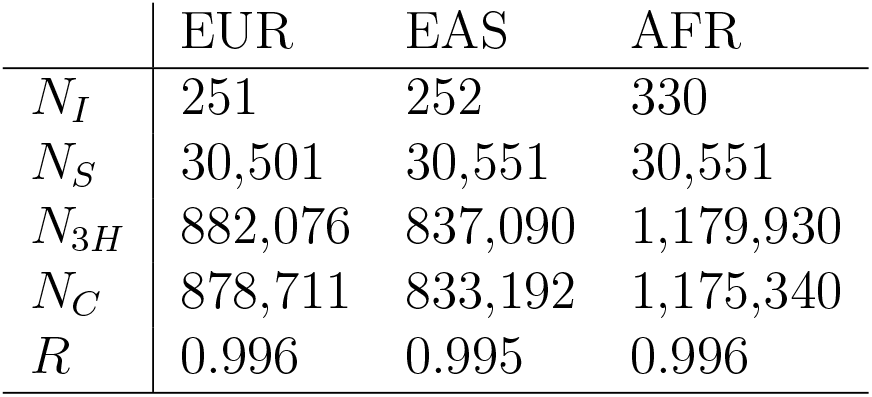
Statistics and accuracy of phasing triple heterozygous sites via mask and imputation. *N*_*i*_: number of simulated samples. *N*_*s*_ number of biallelic SNPs used in simulation. *N*_3*H*_ : number of triple heterozygous SNPs across *N*_*i*_ samples. *N*_*T*_ : number of triple heterozygous SNPs that being correctly phased. *R*: ratio of the correctly phased triple heterozygous SNPs. EUR: European samples. EAS: East Asian samples. AFR: African samples.

## 2 Examples of eGenes harboring different set of eQTL

**Figure S1:**
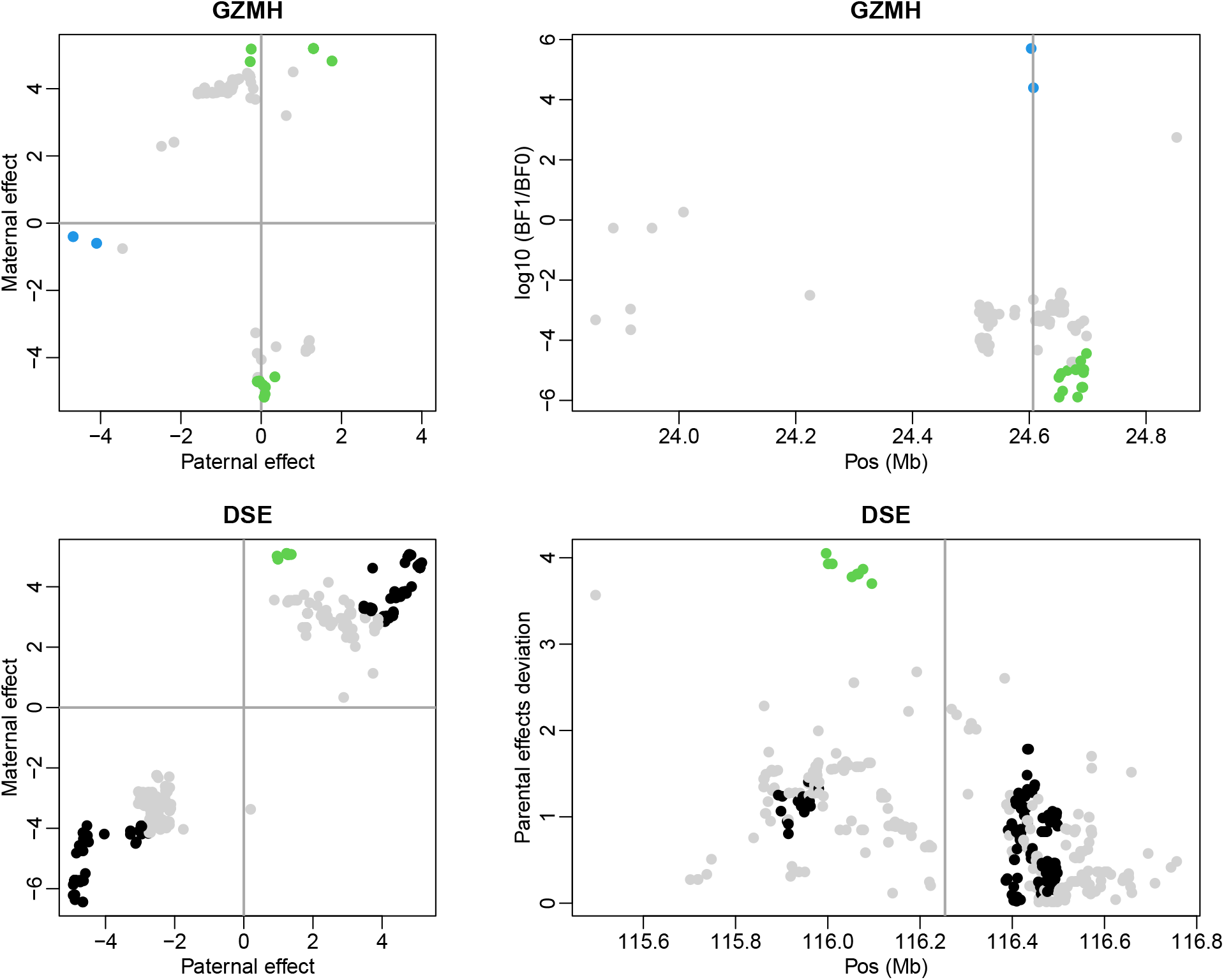
Examples of eGenes harboring different sets of eQTL: *GZMH* and *DSE*. Each gene has two plots: a square plot showing normalized paternal effect (x-axis) vs normalized maternal effect (y-axis), and a rectangle plot showing test statistics along chromosome position. In each plot, gray dots are insignificant eQTL, black dots are significant genotype eQTL, blue dots are significant paternal eQTL, and green dots are significant maternal eQTL. The vertical line in the right panels mark the transcription start site.

## 3 Kinship estimates

**Figure S2:**
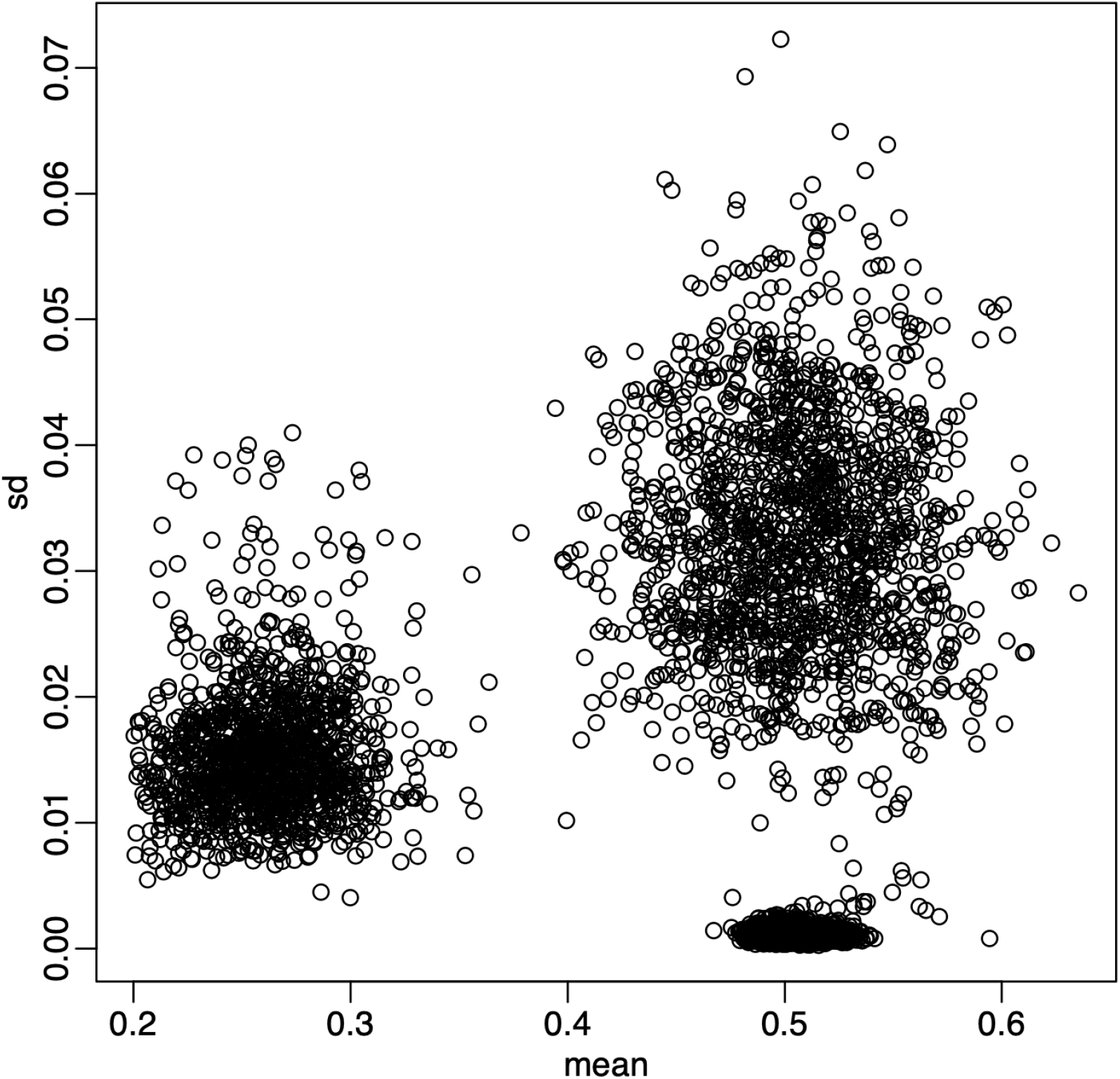
Kinship and sd. The x-axis is twice of kinship averaged over 22 estimates, one for each autosome. The y-axis is standard deviation of those 22 estimates. Plot only show relevant portion of the kinship.

## 4 GC content bias correction

**Figure S3:**
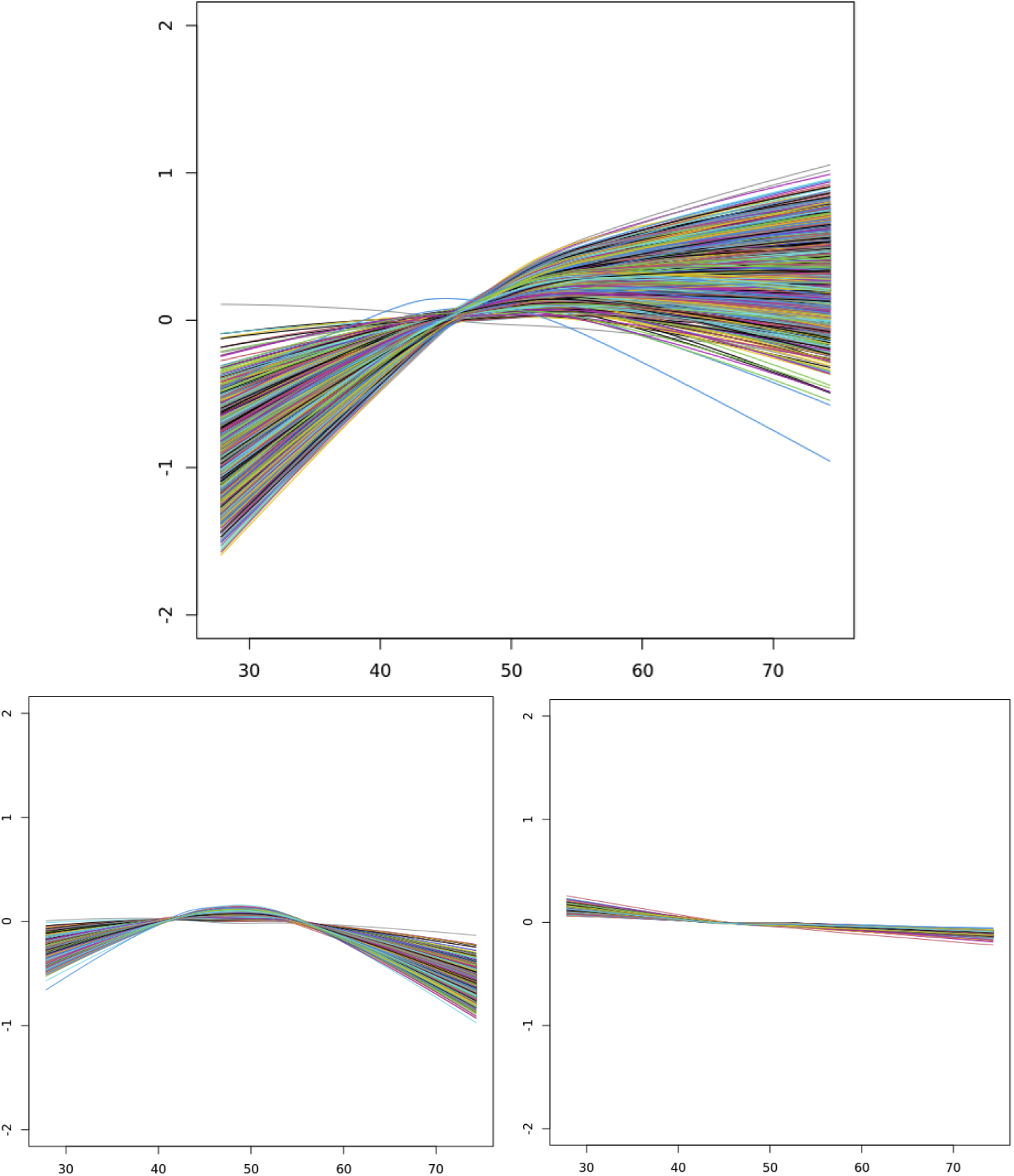
GC content bias correction. Top panel is before correction; Bottom left is after correction with local linear regression; Bottom right is after correction with local quadratic regression (loess in R). Each line represents fitted values of one sample.

## 5 Gene sets for different *θ*

**Table S2:**
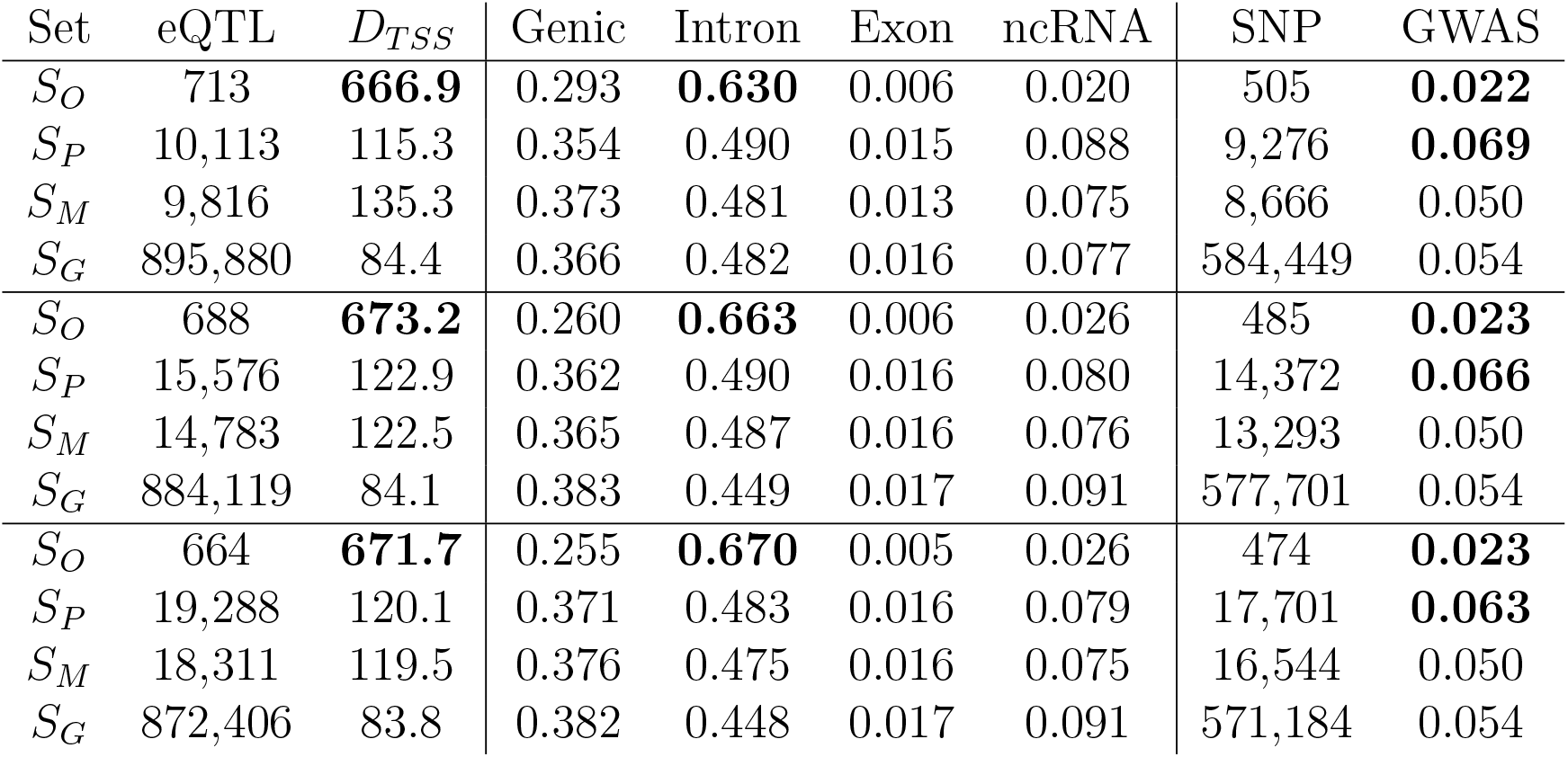
Annotation of eQTL for different threshold. Top tabular: *θ* = 0. Middle tabular *θ* = log_10_ 2. Bottom tabular: threshold= *θ* = log_10_ 3. *S*_*O*_ is a set of opposing eQTL, *S*_*P*_ is a set of paternal eQTL, *S*_*M*_ is a set of maternal eQTL, and *S*_*G*_ is a set of genotype eQTL. The column eQTL contain counts of eQTL in each set, and column SNP contains counts of distinct SNPs of eQTL in the set. The column *eGene* is the number of genes associated with eQTL in the set. The column of Len is the median length in Kb of the eGenes. (The pattern is the same with the mean length.) The column *D*_*TSS*_ contains median distance in Kb to transcription start site. The column GWAS contains percent of GWAS hits among SNPs, with GWAS p-value threshold of 5 *×* 10^−8^.

## 6 Gene sets

### 6.1 Genes harbor exclusively paternal eQTL *G*_1_

*ANGEL1, ANKRD11, ANKS6, AQP3, ARNILA, ASB3, BAG3, BCL2L12, C21orf58, C2orf49-DT, CA11, CAMSAP2, CASP9, CCDC96, CCR9, CDKL3, CEBPZ, CFH, CHKA, CLIP4, CPNE5, CRK, CRY2, DBP, DCANP1, DDX51, DISP2, DLX4, EEF1E1, EHMT1, EPB41L2, FAM136A, FAM220A, FAM50B, FLT3, GEMIN4, GMCL1, GNA15-DT, GRAMD1A, HCLS1, HINFP, HPS6, HSDL1, KPNA2, LCDR, LINC00656, LINC01145, LINC01624, LINC01694, LINC03034, LRIG1, LTBR, MAST2, MBD1, MCEE, MED24, MEF2C-AS1, MICAL2, MIF, MIR22HG, MOCS2, MPHOSPH10, MPP7, MRPL39, NCEH1, NCOA2, NDN, NOPCHAP1, NOTCH2NLC, NPM2, NR1H2, PABPC1, PARP15, PCMTD1-DT, PEG10, PER3, PIAS3, PLPP6, POLD1, PPP1CB-DT, PPP1R14BP3, PRR12, PTGS2, PTK2, RBKS, RNF115, RPL5P1, RRM2, SAGSIN1, SCAF1, SCFD1, SEC14L1P1, SGCE, SLC7A5, SLC9A7P1, SMG1P2, SNRPN, SNURF, SQSTM1, TAF6, TANGO2, TARM1, TCAP, TCEA1P2, TIA1, TIMM23B, TINCR, TMEM129, TMEM30A, TMOD1, TRMT2A, TXNIP, UBR5-DT, UROD, VASH1-AS1, VPS9D1-AS1, WAPL-DT, YWHAZ, ZKSCAN5, ZNF276, ZNF34, ZNF407AS1, ZNF658B, ZNF706, ZNF778-DT, ZNNT1, ZRANB3, ZSCAN16, ZSCAN25*

### 6.2 Genes harbor exclusively maternal eQTL *G*_0_

*ACTBP11, AIP, AKAP8, AMDHD2, ANKRA2, ANKUB1, ANO10, ASAP3, ASCC3, ATG12, BBIP1, BISPR, BLOC1S4, CABP4, CAMKMT, CASS4, CBR3-AS1, CC2D1A, CCDC117, CCDC30, CCT6P1, CDK2AP2, CDKN2AIPNL, CEPT1, CES4A, CMTM3, COA8, CTBP1-DT, CYP2U1, CYSTM1, DDX39A, EIF4B, ELOCP19, EMB, EPS15, EZR, FASTKD3, FEM1A, FUT8-AS1, GALNT4, GLUL, GNL2, GPR108, GRB10, GRK2, GTF2F1, GUCY1B1, GUSB, HAVCR1, HEATR1, HELZ2, HNRNPA1P21, HSPA1A, IGHG4, IGKV2-28, IL23A, ING1, INPP5A,KCTD15, KDM2A, KHSRP, KIR2DL4, LINC00310, LINC00467, LINC02908, LONP1, LRRC8C-DT, LTC4S, LYSMD2, MAN2B2, MAPK8, MCM4, MEG3, METTL15, METTL9, MRFAP1L1, MRPL14 NCF2, NEURL1, NIPSNAP1, NOC4L, NTNG2, NUDT22, OSBPL9, PCM1, PF4, PGGT1B, PI4KAP1, PITPNM1, PLIN3, PMPCB, POC1B-AS1, PPP4R2, PRR5L, PSMB6, PSPN, RANBP3, RASL10A, RCE1, RFX1, RNF43, RPL37AP1, RPS6KB2, RTL10, S1PR2, SCARNA16, SEC23B, SH2D2A, SMIM19, SNHG22, SPRYD7, STAM-DT, STPG3-AS1, SWSAP1, TAGAP, TDP2, TMEM106C, TMEM134, TMEM218, TRAJ37, TRAJ39, TRBV15, UBE2B, UHRF1, USP47, VAV1, VDR, ZBED3-AS1, ZFP37, ZNF286A, ZNF326, ZNF331, ZNF468, ZNF561, ZNF714, ZNF781, ZNF879, ZNF888, ZSWIM4*

### 6.3 Genes harbor exclusively opposing eQTL *G*_2_

*ADAMTSL4-AS1, AFF1, AKR7A2, ANLN, ARID4A, BICD2, BNIP3, BORCS6, BROX, C18orf25, C1orf56, CACNA2D3, CCP110, CD300LF, CD46, CD82, CDK5RAP3, CDKN2A, CENPM, CHD3, CNTROB, COQ7, CPEB4, CPPED1, CXXC5, DNAJB6P1, DPH5-DT, EBLN2, ECSIT, EXOC5, FAM192BP, FBXW7, FGFR1OP2, FXR2, GABARAP, GNB4, GUCY2D, HEBP1, HSCB, IDI1, IQCK, ITPR1, ITPR2, KDM4A, KDM7A, KIAA0586, LEP, LINC01786, LRRC8D-DT, LZIC, MALINC1, MED21, MIR29B2CHG, MPHOSPH8, MRPS14, MZB1, MZT1, NECAB3, NFIA,NNT-AS1, NOTCH3, OPN3, PALLD, PAPOLG, PARPBP, PEX13, PHIP, PIBF1, PIK3CA, PLSCR3, PODNL1, PPP1R3E, PSMA3-AS1, PSMD9, PSTPIP2, PTRHD1, RARA-AS1, RBL2, REL, RNPC3, RPGRIP1L, RPL23P2, RPS28P7, RYBP, SEC61G, SIGLEC15, SLC15A4, STAM, SYNGR2, TMEM182, TMEM40, TOMM6, TPRG1L, UNC45A, WASL, WRAP53, XPO1, ZBTB4, ZDHHC20*

### 6.4 58 phosphoprotein genes

*CCP110, ARID4A, WASL, CHD3, BICD2, AFF1, PSMD9, AKR7A2, XPO1, PSTPIP2, SEC61G, PAPOLG, FGFR1OP2, BORCS6, CDK5RAP3, UNC45A, FBXW7, CACNA2D3, TMEM40, RBL2, PLSCR3, PALLD, PPP1R3E, PHIP, EXOC5, CD46, KDM7A, NOTCH3, NECAB3, ZDHHC20, TPRG1L, C1ORF56, ITPR1, ITPR2, RNPC3, CXXC5, ZBTB4, FXR2, SYNGR2, CD300LF, SLC15A4, KDM4A, OPN3, CDKN2A, BNIP3, STAM, PEX13, CPPED1, ANLN, RYBP, NFIA, KIAA0586, REL, GNB4, CNTROB, MPHOSPH8, CPEB4, WRAP53*

## 7 Sensitivity Analysis

**Table S3:**
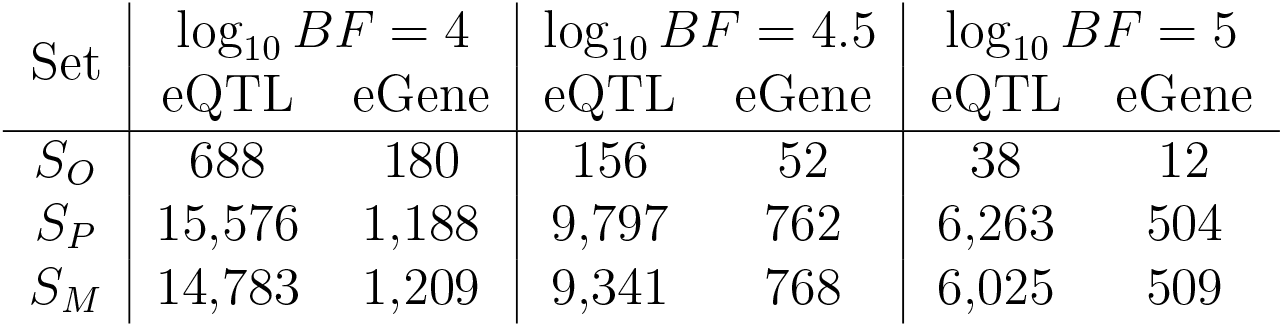
Counts of eQTL and eGenes for different Bayes factor threshold. As the threshold increases, type I error rate reduces, power reduces as a consequence, so the numbers of positive eQTL and eGene reduce.

## 8 Allele Frequencies

**Figure S4:**
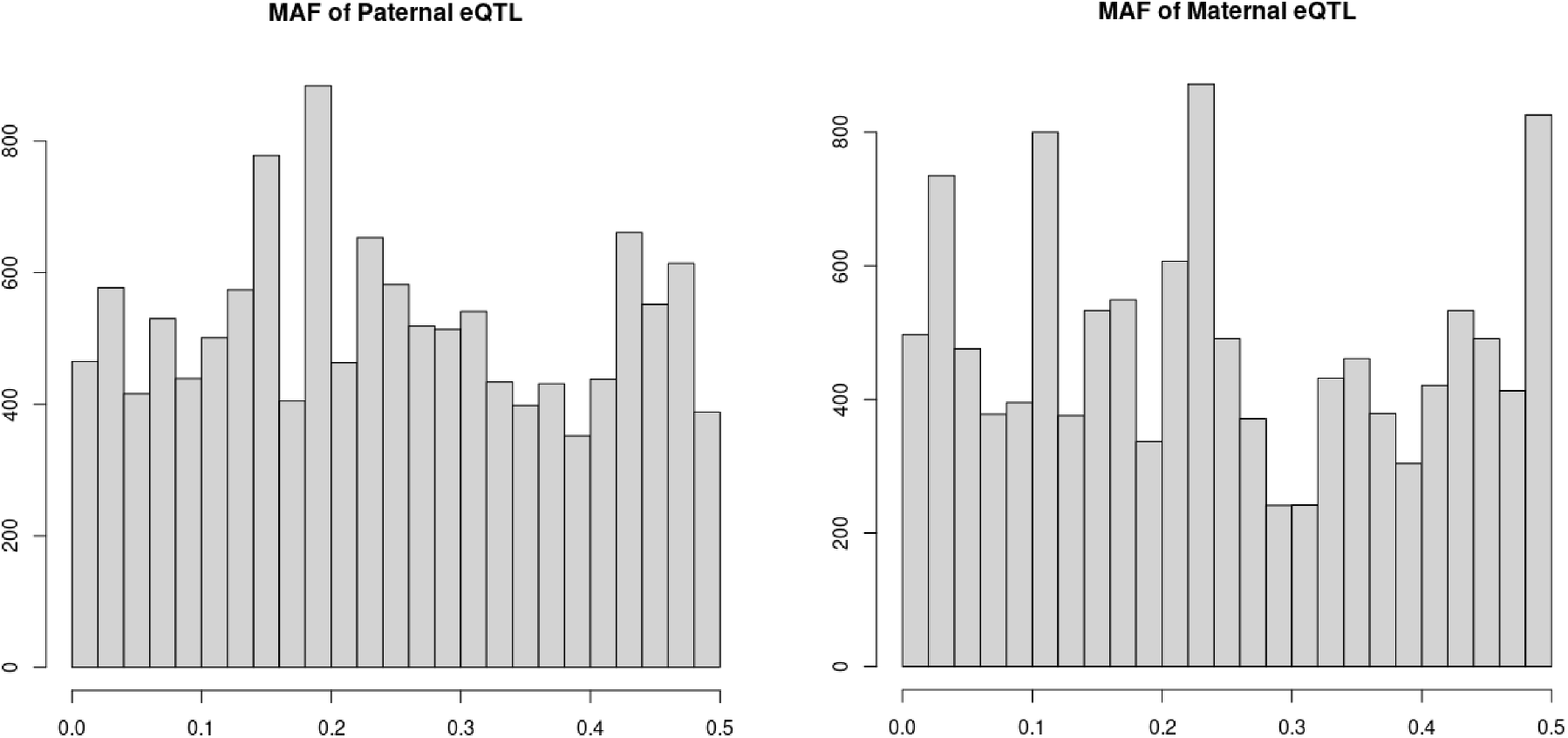
Allele frequecy distribution for parental eQTL and maternal eQTL. Here we used allele frequencies of European descent from 1000 Genomes project.

## 9 P-values, Bayes factors, and FDR

### The p-value and its limitations

By definition, a p-value is the tail probability under the null hypothesis. Formally,

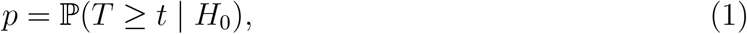

where *T* is a test statistic and *t* = *T*(*D*) is the value computed from the observed data *D*, and *H*_0_ denotes the null hypothesis. It is crucial to emphasize that the p-value is *not* the probability that *H*_0_ is true; it is calculated under the assumption that *H*_0_ holds. In hypothesis testing, we are often interested in the posterior probabilities of the null and alternative hypotheses, ℙ(*H*_0_ | *D*) and ℙ(*H*_1_ | *D*), about which the p-value alone says nothing.

### From p-values to posterior probabilities

Let *R*(*t*) = {*D* : *T*(*D*) ≥ *t*} denote the set of all datasets that would yield a test statistic at least as extreme as the observed value *t*. This *rejection region* extends the observed data to include more extreme unobserved outcomes. Then, by Bayes’ theorem,

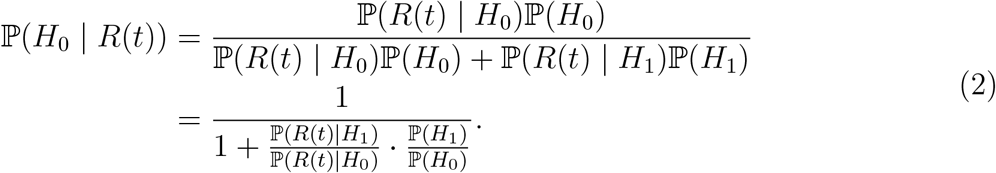

Here, ℙ(*R*(*t*) | *H*_0_) is precisely the p-value, while ℙ(*R*(*t*) | *H*_1_) is the *power* of the test against the alternative hypothesis. The ratio ℙ(*H*_1_)/ℙ(*H*_0_) represents the prior odds. Thus, equation (2) reveals that

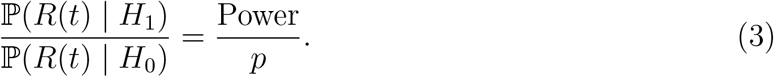

Although a p-value alone conveys no information about posterior probabilities, when combined with power and prior odds, it becomes a useful component for assessing evidence.

### Bayes factors and their relationship to p-values

By definition, the Bayes factor is the ratio of marginal likelihoods under the alternative and null hypotheses:

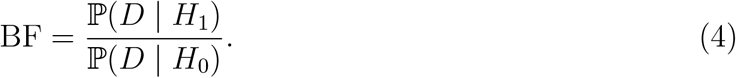

To compute posterior probabilities, one must specify prior odds ℙ(*H*_1_)/ℙ(*H*_0_). The posterior odds are then BF *×* Prior odds, yielding

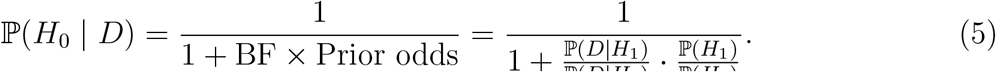

Comparing (2), (3), and (5) suggests an approximate relationship:

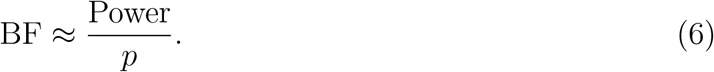

#### Remark 1.

*The approximation in* (6) *is conceptual rather than computational. Power is typically defined at a specific point alternative, whereas* ℙ(*R*(*t*) | *H*_1_) *in* (2) *should properly be integrated over the prior distribution of effect sizes under H*_1_. *This distinction lies at the heart of the Bayesian approach. Nevertheless, this relationship illuminates several important principles:*

- *Interpreting p-values requires consideration of study power*.
- *P-values are not directly comparable across studies with different power*.
- *Underpowered studies require more stringent p-value thresholds to achieve comparable evidence*.

### Local false discovery rate

The local false discovery rate (lfdr) is a Bayesian-inspired measure that assigns a probability of being a false positive to each hypothesis test, conditional on the observed test statistic:

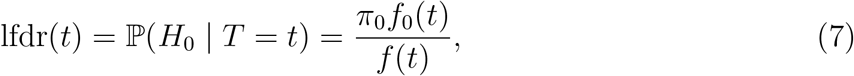

where *f*_0_(*t*) is the density under the null, *f*_1_(*t*) is the density under the alternative, and *f*(*t*) = *π*_0_*f*_0_(*t*)+(1 − *π*_0_)*f*_1_(*t*) is the mixture density. Here, *π*_0_ = ℙ(*H*_0_) is the prior probability of the null. Rearranging,

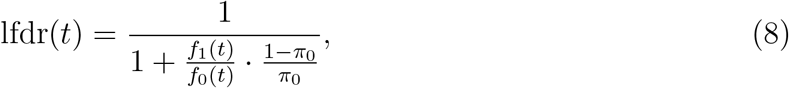

where *f*_1_(*t*)*/f*_0_(*t*) is a Bayes factor (comparing pointwise densities), and (1 − *π*_0_)*/π*_0_ is the prior odds.

### From local fdr to FDR

Let *F*_0_(*t*) = ℙ(*T* ≥ *t* | *H*_0_) be the null tail probability, and *F*_1_(*t*) = ℙ(*T* ≥ *t* | *H*_1_) be the alternative tail probability. The (tail area) false discovery rate at threshold *t* is

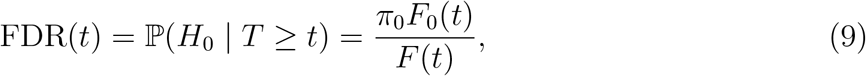

where *F*(*t*) = *π*_0_*F*_0_(*t*) + (1 − *π*_0_)*F*_1_(*t*) is the marginal tail probability. A fundamental connection between local fdr and FDR is:

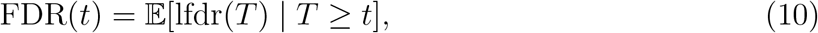

meaning that FDR at threshold *t* is the average local fdr among all tests exceeding that threshold. This relationship illuminates the different philosophical choices: Bayesians often prefer local fdr for its interpretation as an exact conditional probability, while frequentists favor FDR for its reliance on familiar tail probabilities and the convenience of estimating cumulative distribution functions rather than densities.

### The Benjamini-Hochberg procedure as empirical Bayes

The Benjamini-Hochberg (BH) procedure emerges naturally from an empirical Bayes formulation. Let *p*_(1)_ ≤ *p*_(2)_ ≤ *· · ·* ≤ *p*_(*m*)_ be the ordered p-values from *m* hypothesis tests, and let *H*_(*i*)_ denote the null hypothesis corresponding to *p*_(*i*)_. The BH procedure rejects *H*_(1)_, …, *H*_(*k*)_, where *k* is the largest index for which *p*_(*i*)_ ≤ *iα/m*.

To see the connection, consider the empirical estimates of the marginal and null cumulative distribution functions at *p*_(*k*)_:

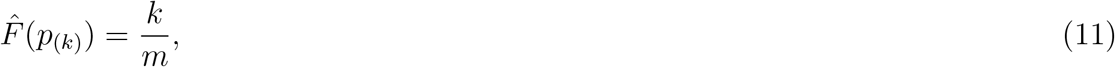

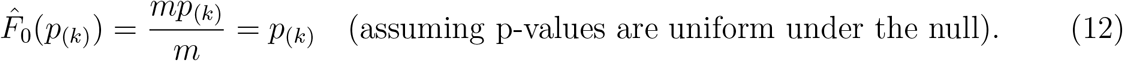

The BH procedure can be derived as rejecting when the estimated FDR falls below *α*:

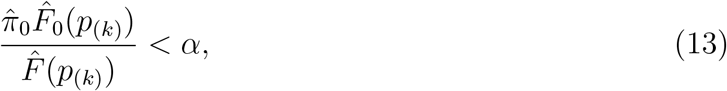

where 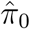 is an estimate of the proportion of true nulls. The original BH procedure implicitly takes 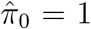 (a conservative choice), yielding *p*_(*k*)_ < *αk/m* as the rejection criterion. More sophisticated procedures estimate *π*_0_ from the data to increase power.

#### Remark 2.

*This derivation reveals that the BH procedure is essentially an empirical Bayes method: it estimates the marginal distribution F empirically through the order statistics, assumes the null distribution F*_0_ *is known (uniform), and rejects tests when the estimated FDR is acceptably low. For cis-eQTL analysis, the uniform null assumption is likely to fail due to LD*.

## Conclusion

P-values, Bayes factors, and FDR-based measures form a coherent framework for understanding statistical evidence. P-values answer a specific probability question under the null, but require context (power and prior odds) to inform posterior beliefs. Its caluation relies on unobserved data, which voilates the likelihood principle. Bayes factors respect the likelihood principle, and provide a direct update of prior odds to posterior odds. The approximate relationship between Bayes factor and p-value in light of its power reflects the advantage of Bayes factor in eQTL analysis. Local fdr and FDR are empirical Bayes procedure in the multiple testing setting, with the BH procedure emerging naturally from an empirical Bayes perspective. Local fdr has an intimate connection with Bayes factor. Understanding these connections helps practitioners avoid common pitfalls in interpreting p-values and choose appropriate methods for their scientific questions.

