## Supplementary for "Abundant Parent-of-origin Effect eQTL: The Framingham Heart Study"

Yongtao Guan, Tianxiao Huan, and Daniel Levy
National Heart, Lung, and Blood Institute

May 3, 2026

#### 1 Simulation study of phasing trios

There are 503 European (EUR), 504 East Asian, and 661 African (AFR) phased samples in 1000 Genomes project. Our simulation was conducted separately for each population. In each population, we randomly selected two samples without replacement and assign the first as father and the second as mother, until exhausted all samples or had only one sample left. Using Chromosome 22, we selected one haplotype from the father and one from the mother to simulate the child. We didn't simulate genotyping error, as in real data the Mendelian error is negligibly small. We then ran the process of phasing by rule of Mendelian inheritance for each trio, masked triple heterozygous SNPs for each child, jointly fit an LD model using all chidren's haplotypes to impute masked SNPs, and finally based on imputed allele dosage to assign alleles (reference or alternative) to maternal and paternal haplotypes. We tallied all counts and put in Table S1, and results show our approach of phasing is highly accurate.

| | EUR | EAS | AFR |
|---|---|---|---|
| $N_I$ | 251 | 252 | 330 |
| $N_S$ | 30,501 | 30,551 | 30,551 |
| $N_{3H}$ | 882,076 | 837,090 | 1,179,930 |
| $N_C$ | 878,711 | 833,192 | 1,175,340 |
| $R$ | 0.996 | 0.995 | 0.996 |

Table S1: Statistics and accuracy of phasing triple heterozygous sites via mask and imputation. $N_i$: number of simulated samples. $N_s$ number of biallelic SNPs used in simulation. $N_{3H}$: number of triple heterozygous SNPs across $N_i$ samples. $N_T$: number of triple heterozygous SNPs that being correctly phased. $R$: ratio of the correctly phased triple heterozygous SNPs. EUR: European samples. EAS: East Asian samples. AFR: African samples.

#### 2 Examples of eGenes harboring different set of eQTL

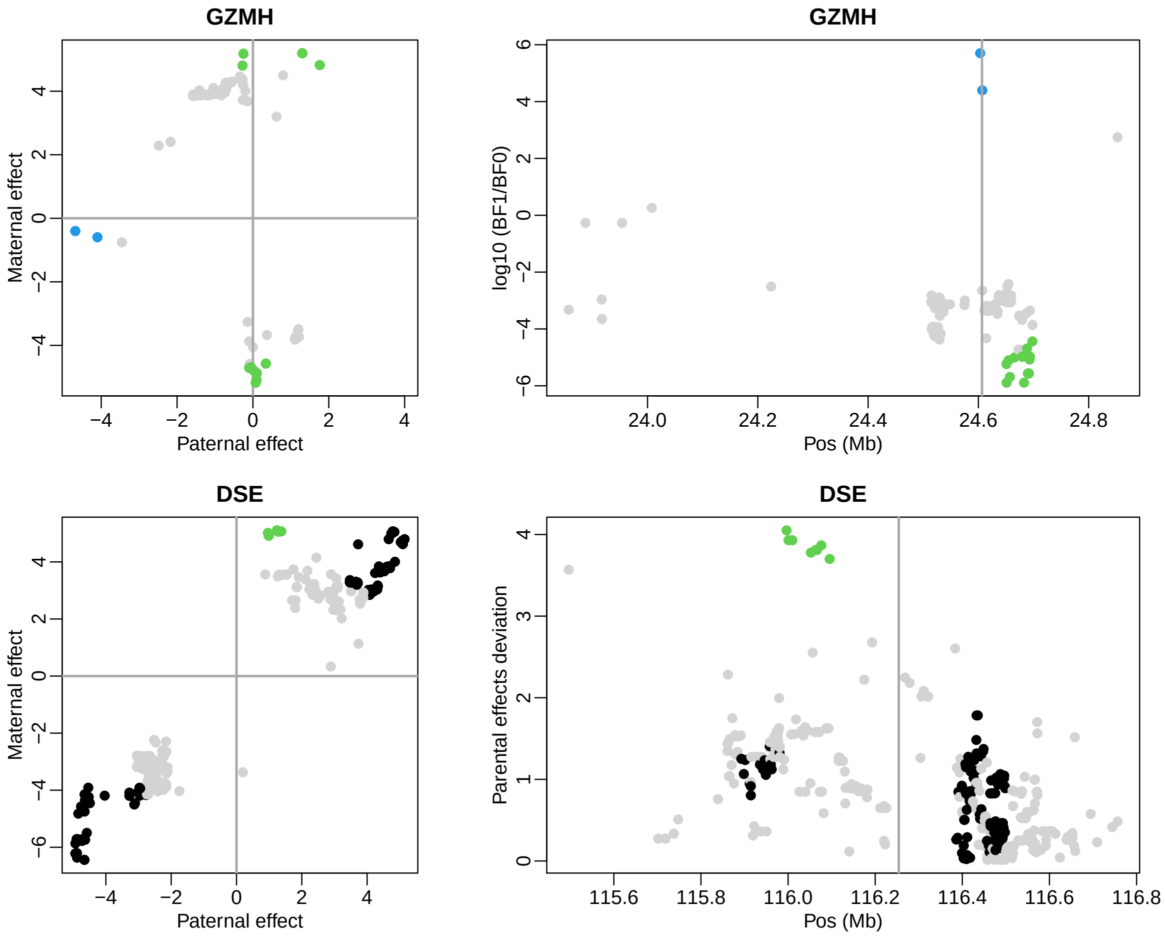


Figure S1: Examples of eGenes harboring different sets of eQTL: *GZMH* and *DSE*. Each gene has two plots: a square plot showing normalized paternal effect (x-axis) vs normalized maternal effect (y-axis), and a rectangle plot showing test statistics along chromosome position. In each plot, gray dots are insignificant eQTL, black dots are significant genotype eQTL, blue dots are significant paternal eQTL, and green dots are significant maternal eQTL. The vertical line in the right panels mark the transcription start site.

### 3 Kinship estimates

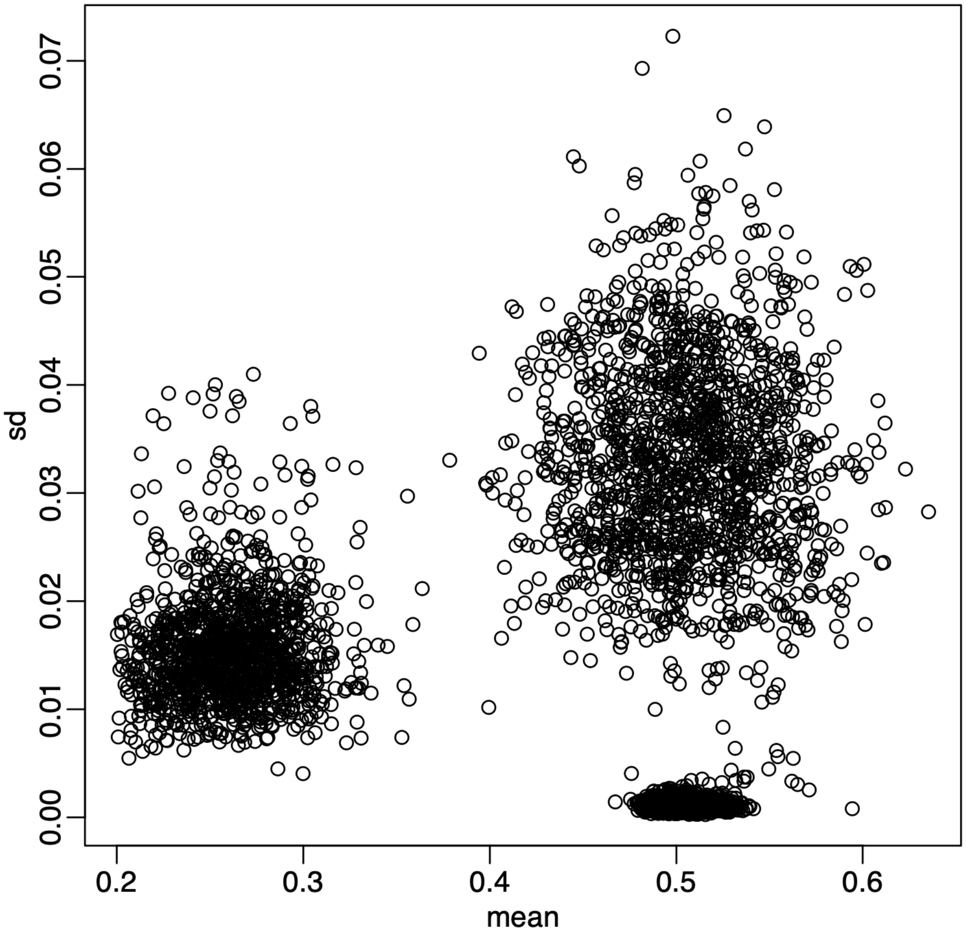


Figure S2: Kinship and sd. The x-axis is twice of kinship averaged over 22 estimates, one for each autosome. The y-axis is standard deviation of those 22 estimates. Plot only show relevant portion of the kinship.

### 4 GC content bias correction

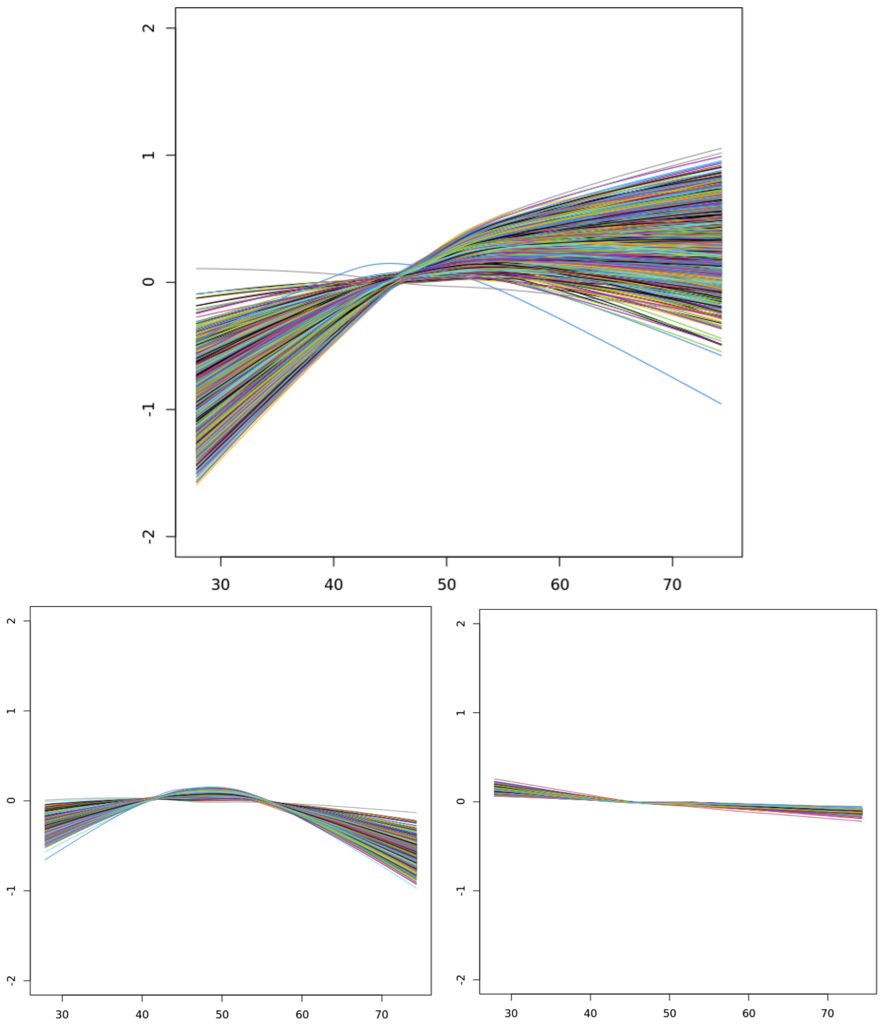


Figure S3: GC content bias correction. Top panel is before correction; Bottom left is after correction with local linear regression; Bottom right is after correction with local quadratic regression (loess in R). Each line represents fitted values of one sample.

### 5 Gene sets for different $\theta$

| Set | eQTL | $D_{TSS}$ | Genic | Intron | Exon | ncRNA | SNP | GWAS |
|---|---|---|---|---|---|---|---|---|
| $S_O$ | 713 | **666.9** | 0.293 | **0.630** | 0.006 | 0.020 | 505 | **0.022** |
| $S_P$ | 10,113 | 115.3 | 0.354 | 0.490 | 0.015 | 0.088 | 9,276 | **0.069** |
| $S_M$ | 9,816 | 135.3 | 0.373 | 0.481 | 0.013 | 0.075 | 8,666 | 0.050 |
| $S_G$ | 895,880 | 84.4 | 0.366 | 0.482 | 0.016 | 0.077 | 584,449 | 0.054 |
| $S_O$ | 688 | **673.2** | 0.260 | **0.663** | 0.006 | 0.026 | 485 | **0.023** |
| $S_P$ | 15,576 | 122.9 | 0.362 | 0.490 | 0.016 | 0.080 | 14,372 | **0.066** |
| $S_M$ | 14,783 | 122.5 | 0.365 | 0.487 | 0.016 | 0.076 | 13,293 | 0.050 |
| $S_G$ | 884,119 | 84.1 | 0.383 | 0.449 | 0.017 | 0.091 | 577,701 | 0.054 |
| $S_O$ | 664 | **671.7** | 0.255 | **0.670** | 0.005 | 0.026 | 474 | **0.023** |
| $S_P$ | 19,288 | 120.1 | 0.371 | 0.483 | 0.016 | 0.079 | 17,701 | **0.063** |
| $S_M$ | 18,311 | 119.5 | 0.376 | 0.475 | 0.016 | 0.075 | 16,544 | 0.050 |
| $S_G$ | 872,406 | 83.8 | 0.382 | 0.448 | 0.017 | 0.091 | 571,184 | 0.054 |

Table S2: Annotation of eQTL for different threshold. Top tabular: $\theta = 0$. Middle tabular $\theta = \log_{10} 2$. Bottom tabular: threshold= $\theta = \log_{10} 3$. $S_O$ is a set of opposing eQTL, $S_P$ is a set of paternal eQTL, $S_M$ is a set of maternal eQTL, and $S_G$ is a set of genotype eQTL. The column eQTL contain counts of eQTL in each set, and column SNP contains counts of distinct SNPs of eQTL in the set. The column *eGene* is the number of genes associated with eQTL in the set. The column of Len is the median length in Kb of the eGenes. (The pattern is the same with the mean length.) The column $D_{TSS}$ contains median distance in Kb to transcription start site. The column GWAS contains percent of GWAS hits among SNPs, with GWAS p-value threshold of $5 \times 10^{-8}$.

#### 7 Sensitivity Analysis

| Set | $\log_{10} BF = 4$ | | $\log_{10} BF = 4.5$ | | $\log_{10} BF = 5$ | |
|---|---|---|---|---|---|---|
| | eQTL | eGene | eQTL | eGene | eQTL | eGene |
| $S_O$ | 688 | 180 | 156 | 52 | 38 | 12 |
| $S_P$ | 15,576 | 1,188 | 9,797 | 762 | 6,263 | 504 |
| $S_M$ | 14,783 | 1,209 | 9,341 | 768 | 6,025 | 509 |

Table S3: Counts of eQTL and eGenes for different Bayes factor threshold. As the threshold increases, type I error rate reduces, power reduces as a consequence, so the numbers of positive eQTL and eGene reduce.

### 8 Allele Frequencies

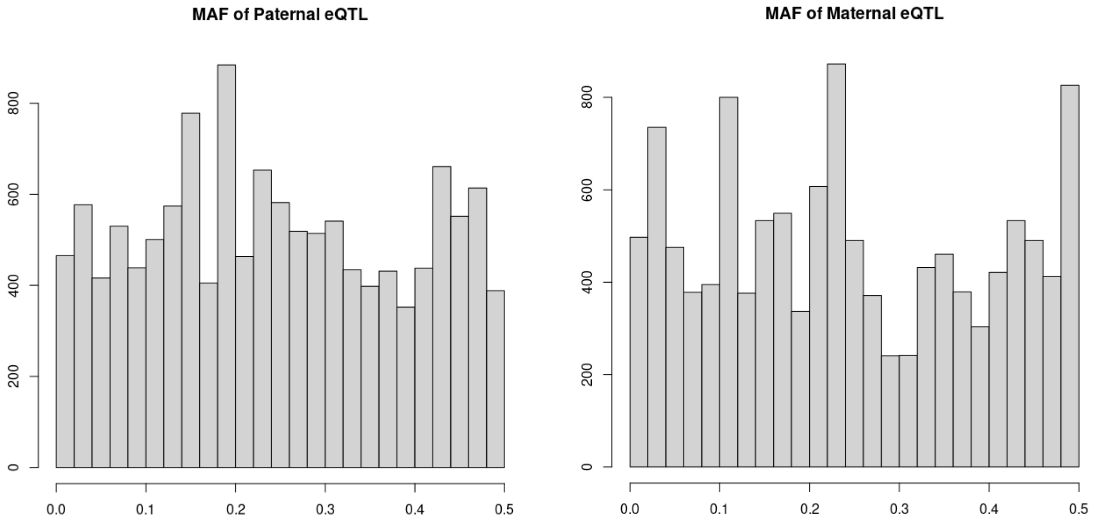


Figure S4: Allele frequecy distribution for parental eQTL and maternal eQTL. Here we used allele frequencies of European descent from 1000 Genomes project.

### 9 P-values, Bayes factors, and FDR

#### The p-value and its limitations

By definition, a p-value is the tail probability under the null hypothesis. Formally,

$$p = \mathbb{P}(T \geq t \mid H_0), \tag{1}$$

where $T$ is a test statistic and $t = T(D)$ is the value computed from the observed data $D$, and $H_0$ denotes the null hypothesis. It is crucial to emphasize that the p-value is *not* the probability that $H_0$ is true; it is calculated under the assumption that $H_0$ holds. In hypothesis testing, we are often interested in the posterior probabilities of the null and alternative hypotheses, $\mathbb{P}(H_0 \mid D)$ and $\mathbb{P}(H_1 \mid D)$, about which the p-value alone says nothing.

#### From p-values to posterior probabilities

Let $R(t) = \{D : T(D) \geq t\}$ denote the set of all datasets that would yield a test statistic at least as extreme as the observed value $t$. This *rejection region* extends the observed data to include more extreme unobserved outcomes. Then, by Bayes' theorem,

$$\begin{aligned} \mathbb{P}(H_0 \mid R(t)) &= \frac{\mathbb{P}(R(t) \mid H_0)\mathbb{P}(H_0)}{\mathbb{P}(R(t) \mid H_0)\mathbb{P}(H_0) + \mathbb{P}(R(t) \mid H_1)\mathbb{P}(H_1)} \\ &= \frac{1}{1 + \frac{\mathbb{P}(R(t)|H_1)}{\mathbb{P}(R(t)|H_0)} \cdot \frac{\mathbb{P}(H_1)}{\mathbb{P}(H_0)}}. \end{aligned} \tag{2}$$

Here, $\mathbb{P}(R(t) \mid H_0)$ is precisely the p-value, while $\mathbb{P}(R(t) \mid H_1)$ is the *power* of the test against the alternative hypothesis. The ratio $\mathbb{P}(H_1)/\mathbb{P}(H_0)$ represents the prior odds. Thus, equation (2) reveals that

$$\frac{\mathbb{P}(R(t) \mid H_1)}{\mathbb{P}(R(t) \mid H_0)} = \frac{\text{Power}}{p}. \tag{3}$$

Although a p-value alone conveys no information about posterior probabilities, when combined with power and prior odds, it becomes a useful component for assessing evidence.

#### Bayes factors and their relationship to p-values

By definition, the Bayes factor is the ratio of marginal likelihoods under the alternative and null hypotheses:

$$\text{BF} = \frac{\mathbb{P}(D \mid H_1)}{\mathbb{P}(D \mid H_0)}. \tag{4}$$

To compute posterior probabilities, one must specify prior odds $\mathbb{P}(H_1)/\mathbb{P}(H_0)$. The posterior odds are then BF $\times$ Prior odds, yielding

$$\mathbb{P}(H_0 \mid D) = \frac{1}{1 + \text{BF} \times \text{Prior odds}} = \frac{1}{1 + \frac{\mathbb{P}(D|H_1)}{\mathbb{P}(D|H_0)} \cdot \frac{\mathbb{P}(H_1)}{\mathbb{P}(H_0)}}. \tag{5}$$

Comparing (2), (3), and (5) suggests an approximate relationship:

$$\mathrm{BF} \approx \frac{\text{Power}}{p}. \tag{6}$$

**Remark 1.** *The approximation in* (6) *is conceptual rather than computational. Power is typically defined at a specific point alternative, whereas* $\mathbb{P}(R(t) \mid H_1)$ *in* (2) *should properly be integrated over the prior distribution of effect sizes under* $H_1$*. This distinction lies at the heart of the Bayesian approach. Nevertheless, this relationship illuminates several important principles:*

- *Interpreting p-values requires consideration of study power.*
- *P-values are not directly comparable across studies with different power.*
- *Underpowered studies require more stringent p-value thresholds to achieve comparable evidence.*

#### Local false discovery rate

The local false discovery rate (lfdr) is a Bayesian-inspired measure that assigns a probability of being a false positive to each hypothesis test, conditional on the observed test statistic:

$$\mathrm{lfdr}(t) = \mathbb{P}(H_0 \mid T = t) = \frac{\pi_0 f_0(t)}{f(t)}, \tag{7}$$

where $f_0(t)$ is the density under the null, $f_1(t)$ is the density under the alternative, and $f(t) = \pi_0 f_0(t) + (1-\pi_0) f_1(t)$ is the mixture density. Here, $\pi_0 = \mathbb{P}(H_0)$ is the prior probability of the null. Rearranging,

$$\mathrm{lfdr}(t) = \frac{1}{1 + \frac{f_1(t)}{f_0(t)} \cdot \frac{1-\pi_0}{\pi_0}}, \tag{8}$$

where $f_1(t)/f_0(t)$ is a Bayes factor (comparing pointwise densities), and $(1 - \pi_0)/\pi_0$ is the prior odds.

#### From local fdr to FDR

Let $F_0(t) = \mathbb{P}(T \geq t \mid H_0)$ be the null tail probability, and $F_1(t) = \mathbb{P}(T \geq t \mid H_1)$ be the alternative tail probability. The (tail area) false discovery rate at threshold $t$ is

$$\mathrm{FDR}(t) = \mathbb{P}(H_0 \mid T \geq t) = \frac{\pi_0 F_0(t)}{F(t)}, \tag{9}$$

where $F(t) = \pi_0 F_0(t) + (1 - \pi_0) F_1(t)$ is the marginal tail probability. A fundamental connection between local fdr and FDR is:

$$\mathrm{FDR}(t) = \mathbb{E}[\mathrm{lfdr}(T) \mid T \geq t], \tag{10}$$

meaning that FDR at threshold $t$ is the average local fdr among all tests exceeding that threshold. This relationship illuminates the different philosophical choices: Bayesians often prefer local fdr for its interpretation as an exact conditional probability, while frequentists favor FDR for its reliance on familiar tail probabilities and the convenience of estimating cumulative distribution functions rather than densities.

#### The Benjamini-Hochberg procedure as empirical Bayes

The Benjamini-Hochberg (BH) procedure emerges naturally from an empirical Bayes formulation. Let $p_{(1)} \leq p_{(2)} \leq \cdots \leq p_{(m)}$ be the ordered p-values from $m$ hypothesis tests, and let $H_{(i)}$ denote the null hypothesis corresponding to $p_{(i)}$. The BH procedure rejects $H_{(1)}, \ldots, H_{(k)}$, where $k$ is the largest index for which $p_{(i)} \leq i\alpha/m$.

To see the connection, consider the empirical estimates of the marginal and null cumulative distribution functions at $p_{(k)}$:

$$\hat{F}(p_{(k)}) = \frac{k}{m}, \tag{11}$$

$$\hat{F}_0(p_{(k)}) = \frac{mp_{(k)}}{m} = p_{(k)} \quad \text{(assuming p-values are uniform under the null).} \tag{12}$$

The BH procedure can be derived as rejecting when the estimated FDR falls below $\alpha$:

$$\frac{\hat{\pi}_0 \hat{F}_0(p_{(k)})}{\hat{F}(p_{(k)})} < \alpha, \tag{13}$$

where $\hat{\pi}_0$ is an estimate of the proportion of true nulls. The original BH procedure implicitly takes $\hat{\pi}_0 = 1$ (a conservative choice), yielding $p_{(k)} < \alpha k/m$ as the rejection criterion. More sophisticated procedures estimate $\pi_0$ from the data to increase power.

**Remark 2.** *This derivation reveals that the BH procedure is essentially an empirical Bayes method: it estimates the marginal distribution $F$ empirically through the order statistics, assumes the null distribution $F_0$ is known (uniform), and rejects tests when the estimated FDR is acceptably low. For cis-eQTL analysis, the uniform null assumption is likely to fail due to LD.*

#### Conclusion

P-values, Bayes factors, and FDR-based measures form a coherent framework for understanding statistical evidence. P-values answer a specific probability question under the null, but require context (power and prior odds) to inform posterior beliefs. Its caluation relies on unobserved data, which voilates the likelihood principle. Bayes factors respect the likelihood principle, and provide a direct update of prior odds to posterior odds. The approximate relationship between Bayes factor and p-value in light of its power reflects the advantage of Bayes factor in eQTL analysis. Local fdr and FDR are empirical Bayes procedure in the multiple testing setting, with the BH procedure emerging naturally from an empirical Bayes perspective. Local fdr has an intimate connection with Bayes factor. Understanding these connections helps practitioners avoid common pitfalls in interpreting p-values and choose appropriate methods for their scientific questions.
